# Database for drug metabolism and comparisons, NICEdrug.ch, aids discovery and design

**DOI:** 10.1101/2020.05.28.120782

**Authors:** Homa MohammadiPeyhani, Anush Chiappino-Pepe, Kiandokht Haddadi, Jasmin Hafner, Noushin Hadadi, Vassily Hatzimanikatis

## Abstract

The discovery of a drug requires over a decade of enormous research and financial investments—and still has a high risk of failure. To reduce this burden, we developed the NICEdrug.ch database, which incorporates 250,000 bio-active molecules, and studied their metabolic targets, fate, and toxicity. NICEdrug.ch includes a unique fingerprint that identifies reactive similarities between drug-drug and drug-metabolite pairs. We use NICEdrug.ch to evaluate inhibition and toxicity by the anticancer drug 5-fluorouracil, and suggest avenues to alleviate its side effects. Clustering based on this fingerprint in statins identified drugs for repurposing. We propose shikimate 3-phosphate for targeting liver-stage malaria with minimal impact on the human host cell. Finally, NICEdrug.ch suggests over 1,300 drugs and food molecules to target COVID-19 and explains their inhibitory mechanisms. The NICEdrug.ch database is accessible online to systematically identify the reactivity of small molecules and druggable enzymes with practical applications in lead discovery and drug repurposing.

## Introduction

To assure effective therapies for previously untreated illness, emerging diseases, and personalized medicine, new small molecules are always needed. However, the process to develop new drugs is complex, costly, and time consuming. This is especially problematic considering about 90% of drug candidates in clinical trials are discarded due to unexpected toxicity or other secondary effects. This inefficiency threatens our health care system and economy (Wong et al., 2019). Improving how we discover and design new drugs could reduce the time and costs involved in the developmental pipeline and hence is of primary importance to define efficient medical therapies.

Current drug discovery techniques often involve high-throughput screens with candidates and a set of target enzymes presumably involved in a disease, which leads to the selection for those candidates with the preferred activity. However, the biochemical space of small molecules and possible targets in the cell is huge, which limits the possible experimental testing. Computational methods for drug pre-screening and discovery are therefore promising. *In silico*, one can systematically search the maximum biochemical space for targets and molecules with desired structures and functions to narrow down the molecules to test experimentally.

There are two main *in silico* strategies for drug discovery: a data-driven approach based on machine learning, or a mechanistic approach based on the available biochemical knowledge. Machine learning (ML) has been successfully used in all stages of drug discovery, from the prediction of targets to the discovery of drug candidates, as shown in some recent studies (Shilo et al., 2020; Stokes et al., 2020; Vamathevan et al., 2019). However, ML approaches require big, high-quality data sets of drug activity and associated physiology (Vamathevan et al., 2019), which might be challenging to obtain when studying drug action mechanisms and side effects in humans. ML also uses trained neural networks, which can lack interpretability and repeatability. This can make it difficult to explain why the neural networks has chosen a specific result, why it unexpectedly failed for an unseen dataset, and the final results may vary (Vamathevan et al., 2019).

Mechanistic-based approaches can also rationally identify small molecules in a desired system and do not require such large amounts of data. Such methods commonly screen based on structural similarity to a native enzyme substrate (antimetabolite) or to a known drug (for drug repurposing), considering the complete structure of a molecule to extract information about protein-ligand fitness (Jarvis and Ouvry, 2019; Verlinde and Hol, 1994). However, respecting enzymatic catalysis, the reactive sites and neighboring atoms play a more important role than the rest of the molecule when assessing molecular reactivity (Hadadi et al., 2019). Indeed, reactive site-centric information might allow to identify: (1) the metabolic fate and neighbors of a small molecule (Javdan et al., 2020), including metabolic precursors or prodrugs and products of metabolic degradation, (2) small molecules sharing reactivity (Lim et al., 2010), and (3) competitively inhibited enzymes (Ghattas et al., 2016). Furthermore, neither ML nor mechanistic-based approaches consider the metabolism of the patient, even though the metabolic fate of the drug and the existence of additional targets in the cell might give rise to toxicity. To our knowledge, no available method accounts for human biochemistry when refining the search for drugs.

In this study, we present the development of the NICEdrug.ch database using a more holistic and updated approach to a traditional mechanistic-based screen by (1) adding a more detailed analysis of drug molecular structures and target enzymes based on structural aspects of enzymatic catalysis and (2) accounting for drug metabolism in the context of human biochemistry. NICEdrug.ch assesses the similarity of the reactivity between a drug candidate and a native substrate of an enzyme based on their common reactive sites and neighboring atoms (i.e., the NICEdrug score) in an analogous fashion as the computational tool BridgIT (Hadadi et al., 2019). It also identifies all biochemical transformations in the cellular metabolism that can modify and degrade a drug candidate using a previously developed reaction prediction tool, termed <u>BI</u>ochemical <u>N</u>etwork <u>I</u>ntegrated <u>C</u>omputational <u>E</u>xplorer (BNICE.ch) (Hatzimanikatis et al., 2005; Soh and Hatzimanikatis, 2010) and the ATLAS of Biochemistry (Hadadi et al., 2016; Hafner et al., 2020). With NICEdrug.ch, we automatically analyzed the functional, reactive, and physicochemical properties of around 250,000 small molecules to suggest the action mechanism, metabolic fate, toxicity, and possibility of drug repurposing for each compound. We apply NICEdrug.ch to study drug action mechanisms and identify drugs for repurposing related to four diseases: cancer, high cholesterol, malaria, and COVID-19. We also sought for molecules in food, as available in fooDB the largest database of food constituents (Scalbert et al., 2011), with putative anti SARS-CoV-2 activity. Finally, we provide NICEdrug.ch as an online resource (https://lcsb-databases.epfl.ch/pathways/Nicedrug/). Overall, NICEdrug.ch combines knowledge of molecular structures, enzymatic reaction mechanisms (as included in BNICE.ch (Finley et al., 2009; Hadadi and Hatzimanikatis, 2015; Hatzimanikatis et al., 2005; Henry et al., 2010; Soh and Hatzimanikatis, 2010; Tokic et al., 2018)), and cellular biochemistry (currently human, *Plasmodium*, and *Escherichia coli* metabolism) to provide a promising and innovative resource to accelerate the discovery and design of novel drugs.

## Results

### Discovery of 200,000 bioactive molecules one reaction away from known drugs in a human cell for analysis of drug metabolism with NICEdrug.ch

To build the initial NICEdrug.ch database, we gathered over 70,000 existing small molecules presumed suitable for treating human diseases from three source databases: KEGG, ChEMBL, and DrugBank (Figure S1, Materials and Methods). We eliminated duplicate molecules, curated available information, computed thermodynamic properties, and applied the Lipinski rules (Lipinski et al., 2001) to keep only the molecules that have drug-like properties in NICEdrug.ch (Figure 1, Materials and Methods). NICEdrug.ch currently includes 48,544 unique small molecules from the source databases.

**Figure 1.**
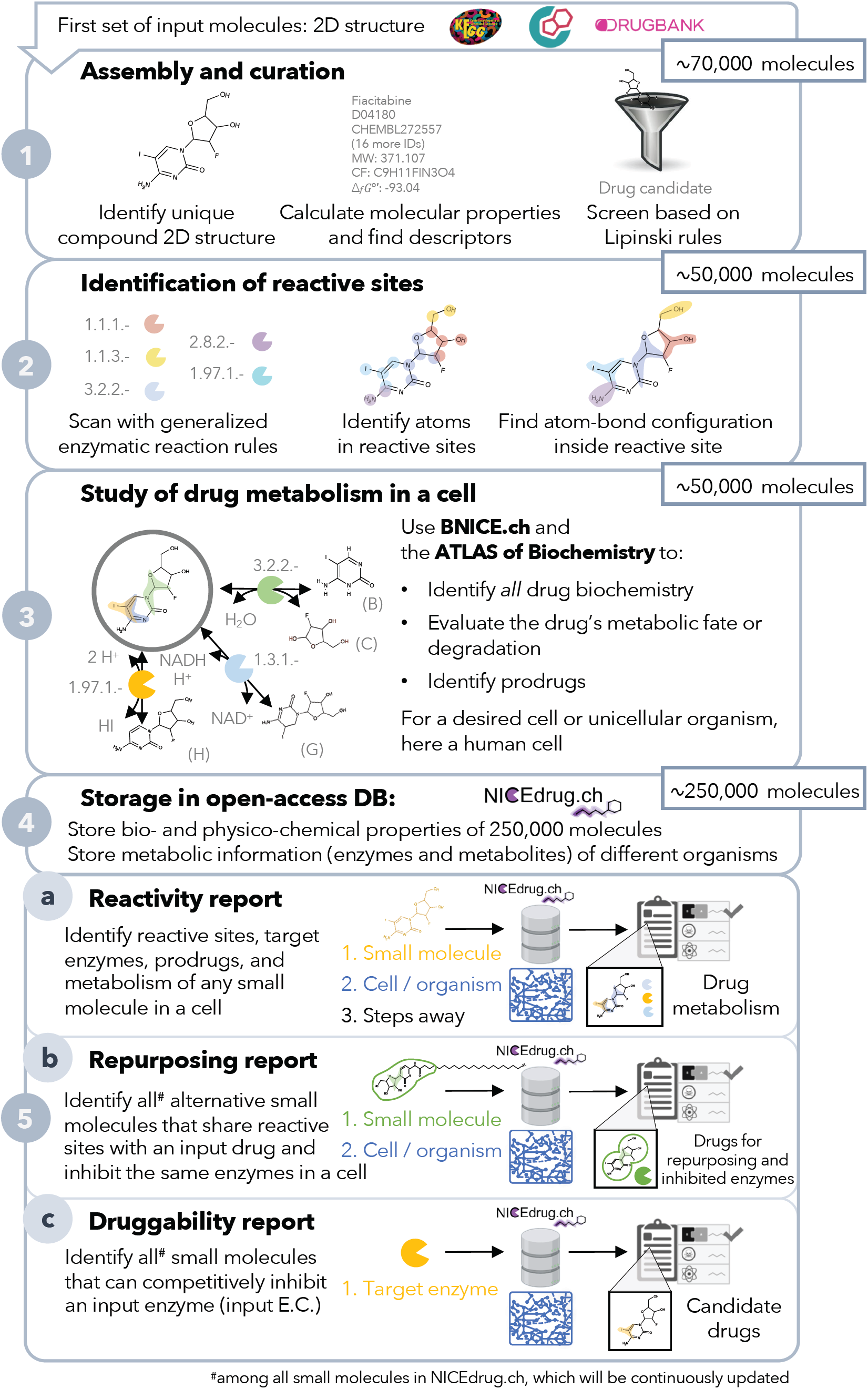
Pipeline to construct and use the NICEdrug.ch database. NICEdrug.ch (1) curates available information and calculates the properties of an input compound; (2) identifies the reactive sites of that compound; (3) explores the hypothetical metabolism of the compound in a cell; (4) stores all functional, reactive, bio-, and physico-chemical properties in open-source database; and (5) allows generation of reports to evaluate (5a) reactivity of a small molecule, (5b) drug repurposing, and (5c) druggability of an enzymatic target. See also Figure S1, Figure S2, Figure S3, and Table S1.

To evaluate the reactivity of the 48,544 drugs, we searched for all possible reactive sites on each drug with BNICE.ch (Hatzimanikatis et al., 2005) (Figure 1, Materials and Methods). All of the 48,544 drugs contain at least one reactive site and hence might be reactive in a cell. In total, we identified more than 5 million potential reactive sites (183k unique) on the 48,544 molecules and matched them to a corresponding enzyme by assigning them to an Enzyme Commission (E.C.) number. All of these enzymes belong to the human metabolic network (Table S1, Materials and Methods). Interestingly, 10.4% of identified reactive sites correspond to the p450 class of enzymes, which are responsible for breaking down compounds in the human body by introducing reactive groups on those compounds, also known as phase I of drug metabolism (Figure S2A). The sites that were identified varied greatly from simple and small (i.e., comprising a minimum number of one atom) to more complex sites that covered a large part of the molecule. The biggest reactive site includes 30 atoms (Figure S2B).

Given the important role of metabolism in the biochemical transformations and toxicity of drugs, we investigated the metabolism of the 48,544 input molecules in human cells. We predicted the hypothetical biochemical neighborhoods of all NICEdrug.ch small molecules in a human cell (i.e., reacting with known human metabolites and cofactors) using a retro-biosynthetic analysis with BNICE.ch (Figure 1, Table S1, Materials and Methods). With this approach, we discovered 197,246 unique compounds connected to the input drugs via one step or reaction (products of the first generation), and the associated hypothetical biochemical neighborhood consists of 630,449 reactions (Figure S2). The 197,246 unique compounds are part of a new set of bioactive molecules in NICEdrug.ch that might act as drugs or prodrugs in a human cell. We stored the total number of 245,790 small molecules (including the curated set of 48,544 drugs and the new set of 197,246 bioactive compounds), their calculated properties, and biochemistry in our open-access database of drug metabolism, NICEdrug.ch.

To use NICEdrug.ch to identify drug-drug or drug-metabolite pairs that have shared reactivity and target enzymes, we developed a new metric called the *NICEdrug score* (Figure S3). The NICEdrug score uses information about the structure of the reactive site and its surroundings (as computed using the BridgIT methodology) and is stored in the form of a fingerprint (Materials and Methods). The fingerprint of a molecule’s reactive site and the neighborhood around this reactive site—termed the *reactive site-centric fingerprint—*serves to compare this site-specific similarity with other molecules. We recently showed that the reactive site-centric fingerprint of a reaction provides a better predictive measure of similar reactivity than the overall molecular structure, as the overall structure can be much larger than the reactive site and skew the results by indicating high similarities when the reactivity is actually quite different (Hadadi et al., 2019). Here, we generated reactive site-centric fingerprints for all 20 million reactive sites identified in the 48,544 drugs and 197,246 one-step-away molecules included in NICEdrug.ch. The 20 million reactive site-centric fingerprints for the total 245,790 small molecules are available in NICEdrug.ch to be used in similarity comparisons and classifying molecules (Materials and Methods).

We propose the usage of NICEdrug.ch to generate reports that define the hypothetical reactivity of a molecule, the molecule’s reactive sites as identified by target enzymes, and the NICEdrug score between drug-drug and drug-metabolite pairs. The NICEdrug.ch reports can be used for three main applications: (1) to identify the metabolism of small molecules; (2) to suggest drug repurposing; and (3) to evaluate the druggability of an enzyme in a desired cell or organism (Figure 1), as we show in the next sections. Currently, NICEdrug.ch includes metabolic information for human cells, a malaria parasite, and *Escherichia coli*, and it is easily extendible to other organisms in the future.

### NICEdrug.ch suggests inhibitory mechanisms of the anticancer drug 5-FU and avenues to alleviate its toxicity

As a case study, we used NICEdrug.ch to investigate the mode of action and metabolic fate of one of the most commonly used drugs to treat cancer, 5-fluorouracil (5-FU), by exploring its reactivity and the downstream products or intermediates that are formed during the cascade of biochemical transformations. 5-FU interferes with DNA synthesis as an antimetabolite (Longley et al., 2003), meaning that its various intermediates like 5-fluorodeoxyuridine monophosphate (FdUMP) are similar enough to naturally occurring substrates and they can act as competitive inhibitors in the cell.

We therefore used NICEdrug.ch to study the intermediates of 5-FU that occurred between one to four reaction steps away from 5-FU (Table S2), which is a reasonable range to occur in the body after 5-FU treatment (Testa, 2010). This analysis identified 407 compounds (90 biochemical and 317 chemical molecules) that have the biochemical potential to inhibit certain enzymes. Because the NICEdrug score that analyses reactive site and neighborhood similarities can serve as a better predictor of metabolite similarity, we assessed the NICEdrug score of the intermediates compared to human metabolites. This resulted in a wide range of NICEdrug scores between the different 5-FU intermediates and human metabolites, ranging from no similarity at a NICEdrug score of 0 to the equivalent substructure on a compound at a NICEdrug score of 1. More importantly, some of the 407 metabolite inhibitors (as explained next) were known compounds that have been investigated for their effects on 5-FU toxicity, but most of these compounds were newly identified by NICEdrug.ch and could therefore serve as avenues for future research into alleviating the side effects of this drug.

We investigated these 407 compounds in more detail, looking first at the set of already validated metabolite inhibitors. 5-Fluorouridine (two steps away from 5-FU) and UDP-L-arabinofuranose (four steps away from 5-FU) are very similar to uridine, with NICEdrug scores of 0.95 and 1, respectively. Uridine is recognized as a substrate by two human enzymes, cytidine deaminase (EC: 3.5.4.5) and 5’-nucleotidase (EC: 3.1.3.5) (Figure 2). Therefore, NICEdrug.ch predictions show that the degradation metabolism of 5-FU generates downstream molecules similar to uridine, which likely leads to the inhibition of these two enzymes. This effect has already been investigated as a potential method for reducing the toxicity of 5-FU, wherein it was proposed that high concentrations of uridine could compete with the toxic 5-FU metabolites (Ma et al., 2017).

**Figure 2.**
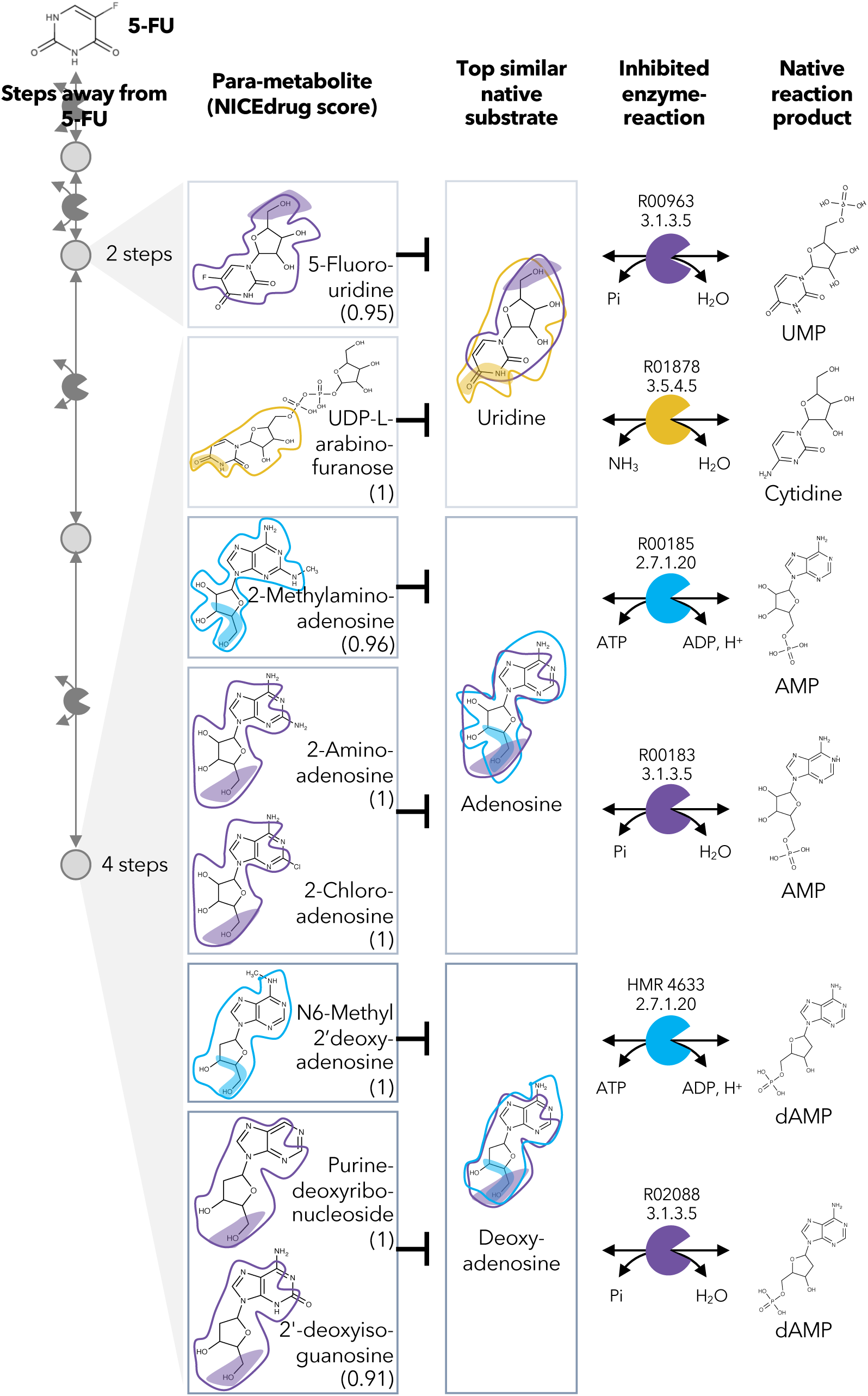
Similarity in reactive site and neighborhood defines para-metabolites in 5-FU metabolism and inhibited human metabolic enzymes. Eight para-metabolites in the 5-FU metabolic neighborhood (represented as defined in Materials and Methods). We show the most similar native human metabolites, inhibited enzymes, and native products of the reactions. See also Table S2.

NICEdrug.ch also identified a few potential metabolites that have not been previously studied for their effects. These metabolites share a reactive site with native human metabolites and differ in the reactive site neighborhood, and we refer to them as *para-metabolites* (Sartorelli and Johns, 2013). 6-Methyl-2’-deoxyadenosine, purine-deoxyribonucleoside, and 2’-deoxyisoguanosine structurally resemble the reactive site neighborhood of deoxyadenosine, with respective NICEdrug scores of 1, 1, and 0.91. Similarly, 2-aminoadenosine, 2-chloroadenosine, and 2-methylaminoadenosine (four steps from 5-FU) have the same reactive site neighborhood as adenosine, with NICEdrug scores of 1, 1, and 0.96, respectively. Adenosine and deoxyadenosine are both native substrates of the adenosine kinase (EC: 2.7.1.20) and 5’-nucleotidase (EC: 3.1.3.5) (Figure 2). Therefore, we suggest that the 5-FU derivatives 2-aminoadenosine and 2-chloroadenosine are competitive inhibitors for the two enzymes adenosine kinase and 5’-nucleotidase. With these new insights from NICEdrug.ch, we hypothesize that co-administering adenosine or deoxyadenosine and uridine (Figure 2) with 5-FU might be required to reduce its toxic effects and hopefully alleviate the side effects of the 5-FU cancer treatment.

### Metabolic degradation of 5-FU leads to compounds with Fluor in their reactive site that are less reactive and more toxic than other intermediates

In the previous case study, we showed inhibitors that contain the identical active site to the native enzyme. However, a slightly different reactive site might still be able to bind to an enzyme and compete with a native substrate, also defined as *anti-metabolite* (Matsuda et al., 2014). We explored this scenario by defining relaxed constraints in two steps. We first identified all atoms around a reactive site to compare the binding characteristics between the native molecule and putative inhibitor. Next, we compared the reactive site of the native molecule and putative inhibitor and scored the latter based on similarity (Materials and Methods). Following these two steps, we assessed the similarity between intermediates in the 5-FU metabolic neighborhood and human metabolites. Among all 407 compounds in the 5-FU metabolism (Table S2), we found 8 that show a close similarity to human metabolites (NICEdrug score above 0.9, Figure 3) that might be competitive inhibitors or anti-metabolites. Inside the reactive site, the original hydrogen atom is bioisosterically replaced by fluorine. F-C bonds are extremely stable and therefore block the active site by forming a stable complex with the enzyme. The inhibitory effect of the intermediates tegafur, 5-fluorodeoxyuridine, and F-dUMP (one to two reaction steps away) has been confirmed in studies by Kobayakawa et.al (Kobayakawa and Kojima, 2011) and Bielas et.al (Bielas et al., 2009). In addition, NICEdrug.ch also predicts that 5flurim, 5-fluorodeoxyuridine triphosphate, 5-fluorodeoxyuridine triphosphate, 5-fluorouridine diphosphate, and 5-fluorouridine triphosphate, some of which occur further downstream in the 5-FU metabolism, also act as antimetabolites (Figure 3). Based on the insights from NICEdrug.ch, we suggest the inhibitory and side effect of 5-FU treatment might be more complex than previously thought. 5-FU downstream products are structurally close to human metabolites and might form stable complexes with native enzymes. This knowledge could serve to further refine the pharmacokinetic and pharmacodynamic models of 5-FU and ultimately the dosage administered during treatment.

**Figure 3.**
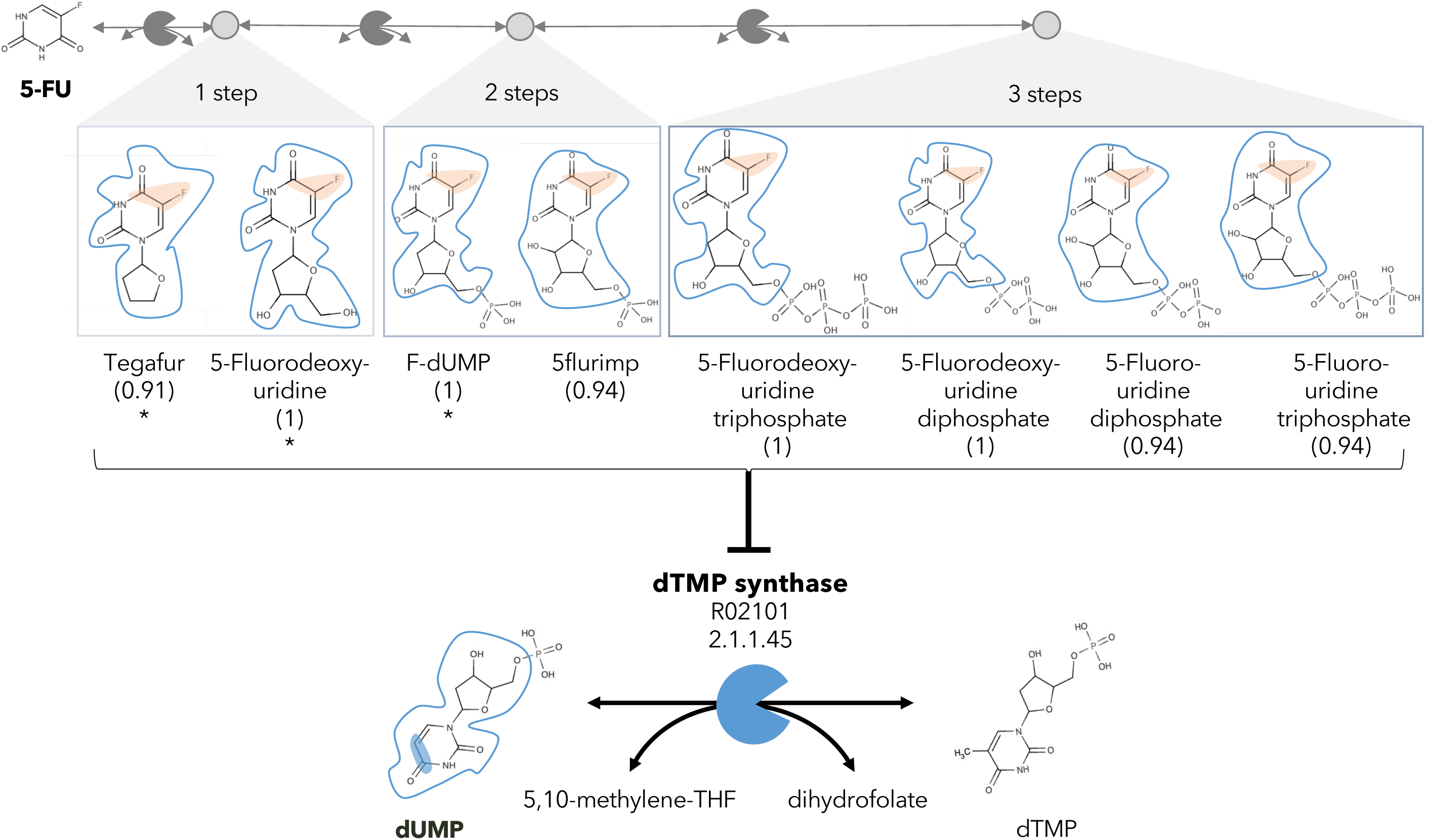
A different reactive site but similar neighborhood defines top anti-metabolites in 5-FU metabolism and inhibited human metabolic enzyme. Eight anti-metabolites of dUMP in the 5-FU metabolic neighborhood (represented as defined in Materials and Methods). Note that the reactive site of the anti-metabolites is different than the one of the native human metabolite, but the neighborhood is highly similar, which determines the high NICEdrug score (value in parenthesis). We show the inhibited human enzyme (dTMP synthase) and reaction, and its native product. See also Table S2.

### NICEdrug.ch identifies toxic alerts in the anticancer drug 5-FU and its products from metabolic degradation

The concept of drug toxicity refers not to overdoses but instead to the toxic effects at medical doses (Guengerich, 2011), which often occur due to the degradation products generated through drug metabolism. Extensive efforts have been expended to identify toxic molecules or, more generally, to extract the substructures that are responsible for toxicity (called structural alerts). The Liver Toxicity Knowledge Base (LTKB) and the super toxic database include 1,036 and about 60k toxic molecules, respectively (Schmidt et al., 2009; Thakkar et al., 2018). ToxAlert provides around 1,200 alerts related to different forms of toxicity (Sushko et al., 2012). However, the number of molecules that are analyzed and labeled as toxic in databases is disproportionally low compared to the space of compounds. Additionally, structural alerts are indicated for many compounds, and current alerts might identify redundant and over-specific substructures, which questions their reliability (Yang et al., 2017).

To quantify the toxicity of downstream products of drugs in NICEdrug.ch, we collected all of the molecules cataloged as toxic in the LTKB and super toxic databases (approved toxic molecules) along with their lethal dose (LC_50_), as well as the existing structural alerts provided by ToxAlert. We measured the similarity of an input molecule with all approved toxic molecules using the reactive site-centric fingerprints implemented in BridgIT and the NICEdrug score (Materials and Methods). Next, we scanned both the toxic reference molecule and the input molecule for structural hints of toxicity, referred to here as *NICEdrug toxic alerts*. We kept common NICEdrug toxic alerts between the reference, which is a confirmed toxic compound, and input molecule. With this procedure in place, NICEdrug.ch finds for each input molecule the most similar toxic molecules along with their common toxic alerts and serves to assess the toxicity of a new molecule based on the mapped toxic alerts. Additionally, the NICEdrug toxic alerts and toxicity level of drug intermediates can be traced with NICEdrug.ch through the whole degradation pathway to reveal the origin of the toxicity.

As an example, we herein tested the ability of NICEdrug.ch to identify the toxicity in 5-FU metabolism. First, we queried the toxicity profile of all intermediates in the 5-FU metabolic neighborhood, integrating both known and hypothetical human reactions (Materials and Methods). In this analysis, we generated all compounds up to four steps away from 5-FU. Based on the toxicity report of each potential degradation product, we calculated a relative toxicity metric that adds the LC_50_ value, NICEdrug score, and number of common NICEdrug toxic alerts with all approved toxic drugs (Materials and Methods). We generated the metabolic neighborhood around 5-FU, and labeled each compound with our toxicity metric (Table S2). Interestingly, we show that the top most toxic intermediates match the list of known three toxic intermediates in 5-FU metabolism (Figure 4) (Krauß and Bracher, 2018). Based on the toxicity analysis in NICEdrug.ch for 5-FU, we hypothesize there are highly toxic products of 5-FU drug metabolism that had not been identified either experimentally or computationally and it might be necessary to experimentally evaluate their toxicity to recalibrate the dosage of 5-FU treatment.

**Figure 4.**
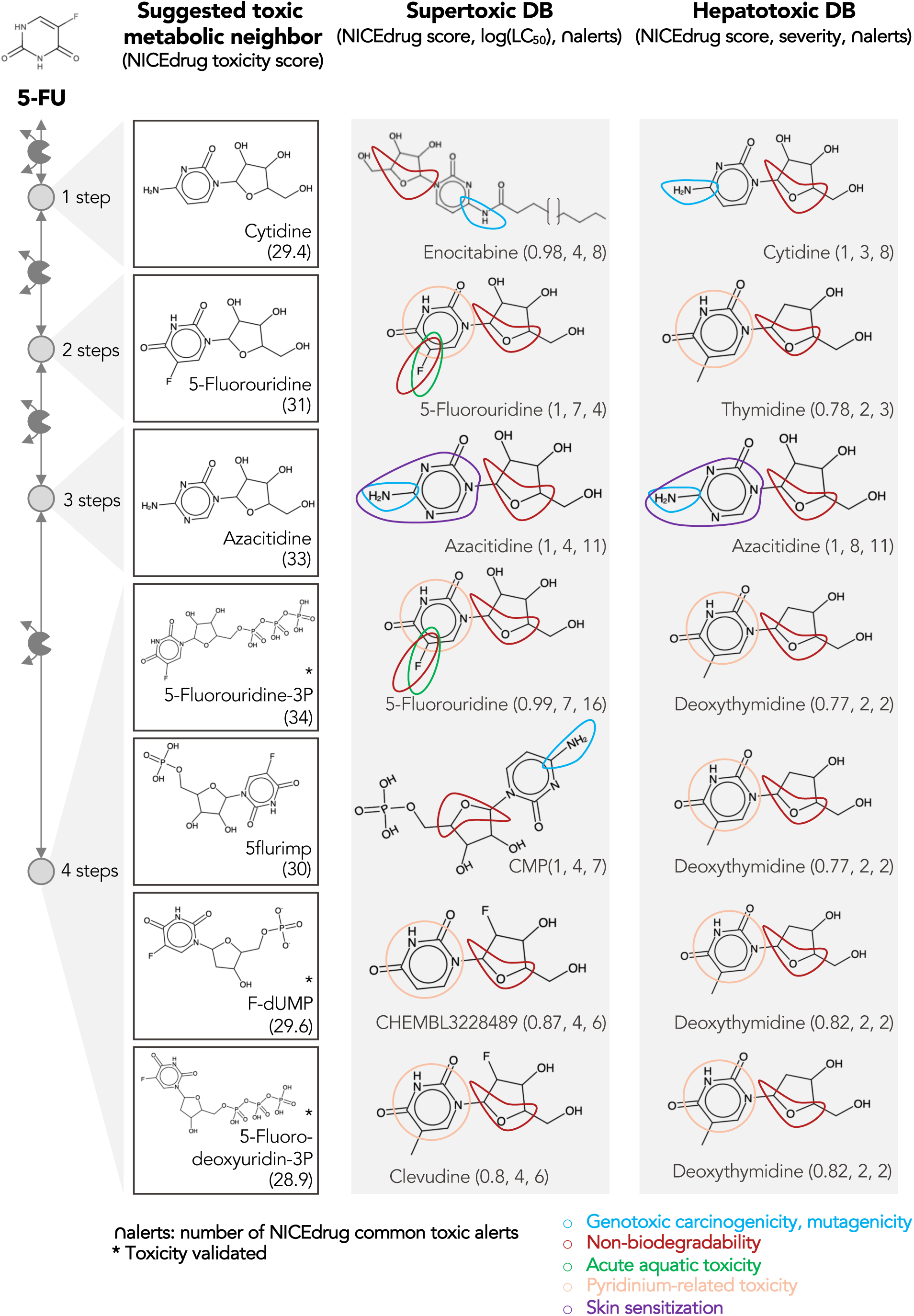
Comparing downstream products to known toxic molecules and analyzing their common structural toxic alerts explains metabolic toxicity of 5-FU. Example of six suggested toxic molecules in the 5-FU metabolic neighborhood (represented as defined in Materials and Methods). We show toxic compounds from the supertoxic and hepatotoxic databases that lead to the highest NICEdrug toxicity score (number under toxic intermediate name, Materials and Methods). We highlight functional groups linked to five NICEdrug toxic alerts (legend bottom right). See also Table S2.

### The NICEdrug reactive site-centric fingerprint accurately clusters statins of type I and II and guides drug repurposing

Because potential side effects of a drug are documented when the drug passes the approval process, repurposing approved drugs for other diseases can reduce the medical risks and development expenses. For instance, the antitussive noscapine has been repurposed to treat some cancers (Mahmoudian and Rahimi-Moghaddam, 2009; Rajesh, A. and International, 2011). Because NICEdrug.ch can search for functional (i.e., reactivity), structural (i.e., size), and physicochemical (i.e., solubility) similarities between molecules while accounting for human biochemistry, we wanted to determine if NICEdrug.ch could therefore suggest drug repurposing strategies.

As a case study, we investigated the possibility of drug repurposing to replace statins, which are a class of drugs often prescribed to lower blood cholesterol levels and to treat cardiovascular disease. Indeed, data from the National Health and Nutrition Examination Survey indicate that nearly half of adults 75 years and older in the United States use prescription cholesterol-lowering statins (US Preventive Services Task Force, 2016). Since some patients do not tolerate these drugs and many still do not reach a safe blood cholesterol level (Kong et al., 2004), there is a need for alternatives. Being competitive inhibitors of the cholesterol biosynthesis enzyme 3-hydroxy-3-methyl-glutaryl-coenzyme A reductase (HMG-CoA reductase) (Jiang et al., 2018; Mulhaupt et al., 2003), all statins share the same reactive site. BNICE.ch labeled this reactive site, in a linear or circular form, as corresponding to an EC number of 4.2.1.- (Istvan, 2001). NICEdrug.ch includes 254 molecules with the same reactive site that are recognized by enzymes of E.C. class 4.2.1.-, ten of which are known statins. We used the NICEdrug score to cluster the 254 molecules into different classes (Table S3, Figure 5). Two of the classes correspond to all currently known statins, which are classified based on their activity into type 1 and 2, wherein statins of type 2 are less active and their reactive site is more stable compared to type 1. This property is well distinguished in the clustering based on the NICEdrug score (Figure 5A).

**Figure 5.**
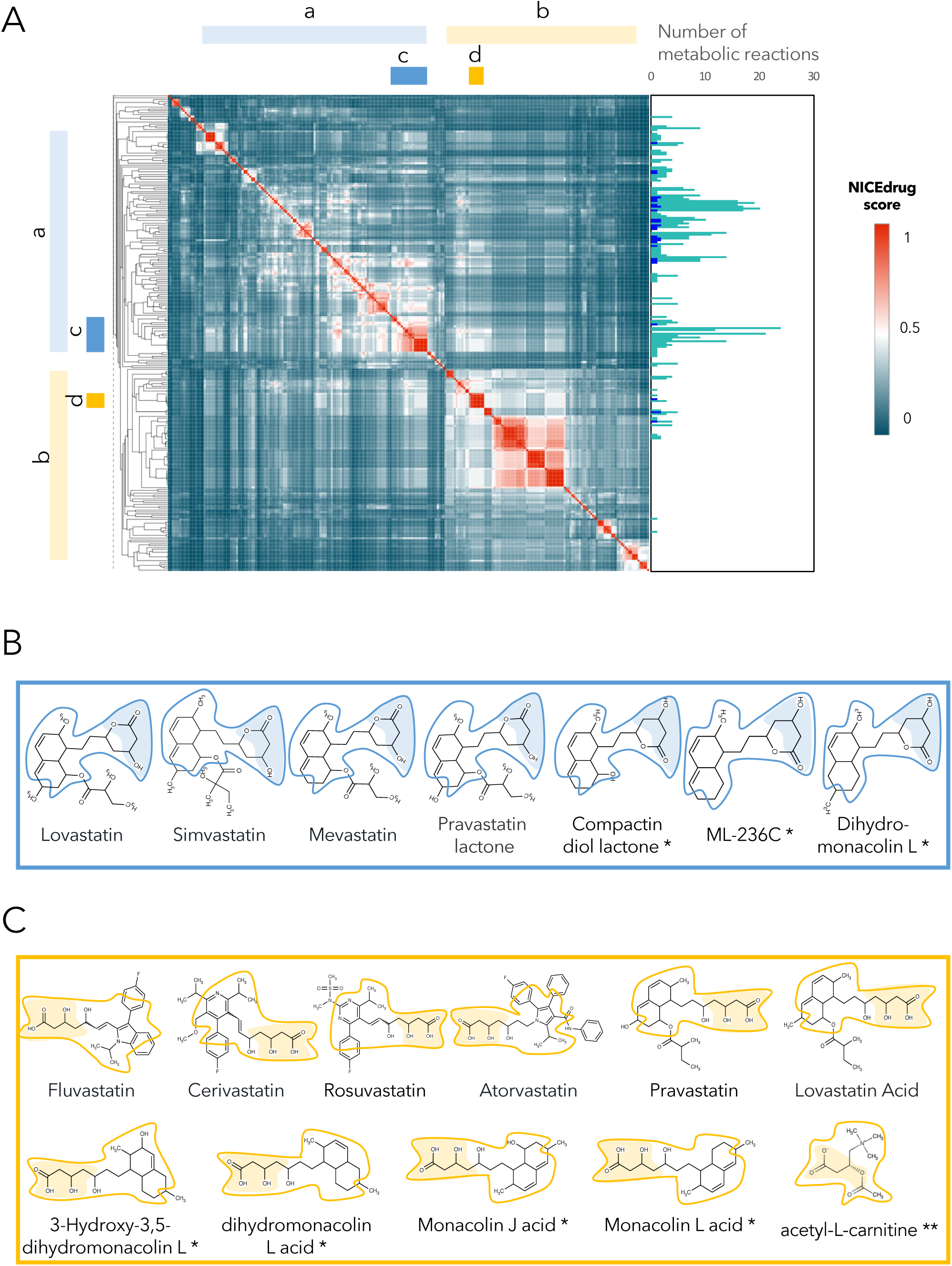
Clustering of molecules with statin reactive sites based on NICEdrug score suggests drugs for repurposing. **(A)** Pairwise NICEdrug score between all molecules with statin reactive sites (heat map) and number of metabolic reactions in which they participate (right). We highlight clusters of statins of type 1 (cluster a) and type 2 (cluster b), and clusters of most similar molecules to type 1 statins (cluster c) and type 2 statins (cluster d). Within the metabolic reactions, we indicate the total number of reactions (dark color) and the number of reactions that involve the statin reactive site (light color). **(B)** Examples of statins and Mevastatin analogues of type 1 from cluster c (blue) and of type 2 from cluster d (gold). We left the known statins unmarked, which are appropriately clustered together based on the NICEdrug score, and we mark with * new molecules that cluster with statins and that NICEdrug.ch suggests could be repurposed to act as statins. Reactive sites in type 1 statins and type 2 statins are colored in blue and orange, respectively. The reactive site neighborhood as considered in the NICEdrug score is also marked. See also Figure S3, Figure S4, Table S3, and Table S4.

In addition to properly classifying the ten known statins (Figure 5B and 5C, molecules non-marked), we identified seven other NICEdrug.ch molecules that clustered tightly with these statins (Figure 5B and 5C, molecules marked with *). These new molecules share the same reactive site and physicochemical properties, and they have the highest similarity with known statins in atoms neighboring the reactive site. In a previous study by Endo *et al.*, these seven NICEdrug.ch molecules were introduced as Mevastatin analogues for inhibiting cholesterol biosynthesis (Endo and Hasumi, 1993). Therefore, they were already suggested as possible candidates for treating high blood cholesterol and could be a good option for repurposing. Furthermore, we found eight known drugs not from the statin family among the 254 scanned molecules (Table S4). One of them, acetyl-L-carnitine (Figure 5C, molecule marked with **), is mainly used for treating neuropathic pain (Li et al., 2015), though Tanaka *et al.* have already confirmed that it also has a cholesterol-reducing effect (Tanaka et al., 2004).

Overall, NICEdrug.ch was able to characterize all known enzymatic reactions that metabolize statins, including proposed alternatives and new hypothetical reactions that could be involved in their metabolism within human cells (Figure 5A, Figure S4). The identification of seven drugs that clustered around the statins and were already designed as alternatives to statins verifies the ability of NICEdrug.ch and the NICEdrug score to search broad databases for similar compounds in structure and function. Furthermore, the discovery of the eight compounds unrelated to known statins offer multiple candidate repurposable drugs along with a map of their metabolized intermediates for the treatment of high cholesterol, though further preclinical experiments would be required to verify their clinical benefits.

### NICEdrug.ch suggests over 500 drugs to target liver-stage malaria and simultaneously minimize side effects in human cells, with shikimate 3-phosphate as a top candidate

Efficiently targeting malaria remains a global health challenge. Malaria parasites (*Plasmodium*) are developing resistance to all known drugs, and antimalarials cause many side effects (World Health Organization, 2018). We applied NICEdrug.ch to identify drug candidates that target liver-stage developing malaria parasites and lessen or avoid side effects in human cells.

We previously reported 178 essential genes and enzymes for liver-stage development in the malaria parasite *Plasmodium berghei* (Stanway et al., 2019) (Table S5, STAR Methods). Out of 178 essential *Plasmodium* enzymes, 32 enzymes are not essential in human cells (Wang et al., 2015) (Table S5, STAR Methods). We extracted all molecules catalyzed by these 32 enzymes uniquely essential in *Plasmodium*, which resulted in 68 metabolites and 157 unique metabolite-enzyme pairs (Table S5, STAR Methods). We used NICEdrug.ch to examine the druggability of the 32 essential *Plasmodium* enzymes with the curated 48,544 drugs (Figure 1) and the possibility of repurposing them to target malaria.

We considered as candidates for targeting liver-stage malaria as the drugs or their metabolic neighbors that show a good NICEdrug score (NICEdrug score above 0.5) with any of the 157 *Plasmodium* metabolite-enzyme pairs. We identified 516 such drug candidates, targeting 16 essential *Plasmodium* enzymes (Table S6, STAR Methods). Furthermore, 1,164 other drugs appear in the metabolic neighborhood of the 516 identified drugs (between one and three reaction steps away). Interestingly, out of the 516 identified drug candidates, digoxigenin, estradiol-17beta and estriol have been previously validated as antimalarials (Antonova-Koch et al., 2018) and NICEdrug.ch suggests their antimalarial activity relies on the competitive inhibition of the KRC enzyme (Figure 6). This enzyme is part of both the steroid metabolism and the fatty acid elongation metabolism, which we recently showed is essential for *Plasmodium* liver-stage development (Stanway et al., 2019). Among the 516 NICEdrug antimalarial candidates, there are also 89 molecules present in the metabolic neighborhood of antimalarial drugs approved by (Antonova-Koch et al., 2018), which suggests these antimalarials might be prodrugs (Table S6).

**Figure 6.**
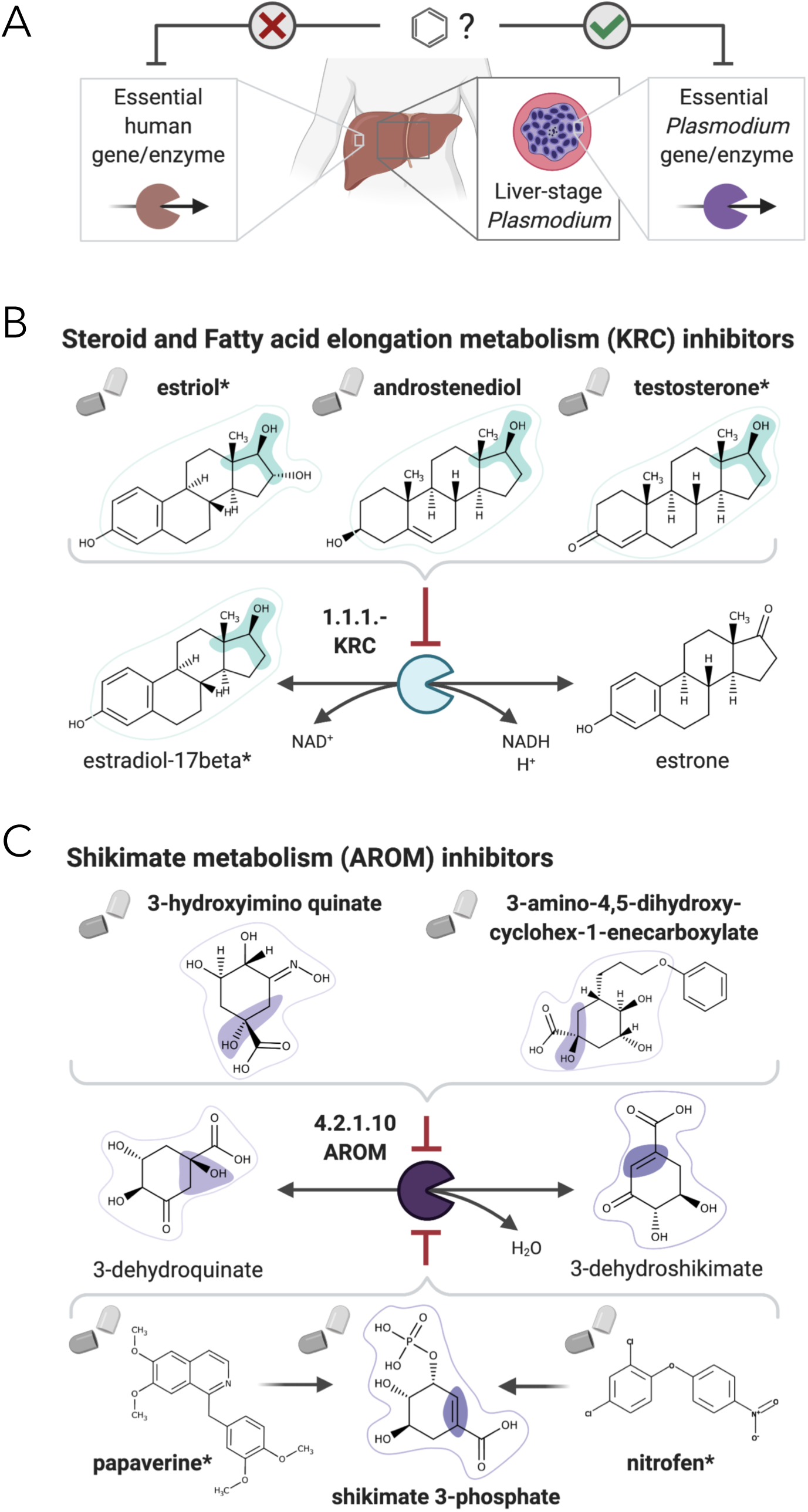
NICEdrug.ch suggests shikimate 3-phosphate as a top candidate to target liver-stage malaria and minimize side effects in host human cells. **(A)** Schema of ideal scenario to target malaria, wherein a drug efficiently inhibits an essential enzyme for malaria parasite survival and does not inhibit essential enzymes in the host human cell to prevent side effects. **(B)** Shikimate 3-phosphate inhibits enzymes in the *Plasmodium* shikimate metabolism, which is essential for liver-stage development of the parasite. Shikimate 3-phosphate does not inhibit any enzyme in the human host cell since it is not a native human metabolite, and it does not show similarity to any native human metabolite. **(C)** Mechanistic details of inhibition of aroC by shikimate 3-phosphate and other NICEdrug candidates. See also Table S5 and Table S6.

Being an intracellular parasite, antimalarial treatments should be efficient at targeting *Plasmodium* as well as assure the integrity of the host cell (Figure 6A). To tackle this challenge, we identified 1,497 metabolites participating in metabolic reactions catalyzed with essential human enzymes (Table S5, STAR Methods) and excluded the antimalarial drug candidates that shared reactive site-centric similarity with the extracted human metabolite set (to satisfy NICEdrug score below 0.5). Out of all 516 drug candidates that might target liver-stage *Plasmodium*, a reduced set of 64 molecules minimize the inhibition of essential human enzymes (Table S6, STAR Methods) and are hence optimal antimalarial candidates.

Among our set of 64 optimal antimalarial candidates, a set of 14 drugs targeting the *Plasmodium* shikimate metabolism, whose function is essential for liver-stage malaria development (Stanway et al., 2019), arose as the top candidate because of its complete absence in human cells. The set of drugs targeting shikimate metabolism include 40 prodrugs (between one and three reaction steps away) that have been shown to have antimalarial activity (Antonova-Koch et al., 2018) (Table S6). NICEdrug.ch identified molecules among the prodrugs with a high number of toxic alerts, like nitrofen. It also identified four molecules with scaffolds similar (two or three steps away) to the 1-(4-chlorobenzoyl)pyrazolidin-3-one of shikimate and derivatives. This result suggests that downstream compounds of the 40 prodrugs might target the *Plasmodium* shikimate pathway, but also might cause side effects in humans (Table S6).

To this end, NICEdrug.ch identified shikimate 3-phosphate as a top candidate antimalarial drug. We propose that shikimate 3-phosphate inhibits the essential *Plasmodium* shikimate biosynthesis pathway without side effects in the host cell (Figure 6, Table S6). Excitingly, shikimate 3-phosphate has been used to treat *E. coli* and *Streptococcus* infections without appreciable toxicity for patients (Díaz-Quiroz et al., 2018). Furthermore, recent studies have shown that inhibiting the shikimate pathway using 7-deoxy-sedoheptulose is an attractive antimicrobial and herbicidal strategy with no cytotoxic effects on mammalian cells (Brilisauer et al., 2019). Experimental studies should now validate the capability of shikimate 3-phosphate to efficiently and safely target liver malaria, and could further test other NICEdrug.ch antimalarial candidates (Table S6).

### NICEdrug.ch identifies over 1,300 molecules to fight COVID-19, with N-acetylcysteine as a top candidate

SARS-CoV-2 is responsible for the currently on-going COVID-19 pandemic and the death of around half a million people (as of today, June 15 (Dong et al., 2020)) and there is currently no confirmed treatment for it. Attacking the host factors that allow replication and spread of the virus is an attractive strategy to treat viral infections like COVID-19. A recent study has identified 332 interactions between SARS-CoV-2 proteins and human proteins, which involve 332 hijacked human proteins or host factors (Gordon et al., 2020). Here, we first used NICEdrug.ch to identify inhibitors of enzymatic host factors of SARS-CoV-2. Targeting such human enzymes prevents interactions between human and viral proteins (PPI) (STAR Methods, Figure 7A). Out of the 332 hijacked human proteins we identified 97 enzymes (STAR Methods, Table S7) and evaluated their druggability by inhibitors among the 250,000 small molecules in NICEdrug.ch and 80,000 molecules in food (STAR Methods, Figure 7A). NICEdrug.ch suggests 22 hijacked human enzymes can be drug targets, and proposed 1301 potential competitive inhibitors from the NICEdrug.ch database. Out of 1301 potential inhibitors, 465 are known drugs, 712 are active metabolic products of 1,419 one-step-away prodrugs, and 402 are molecules in fooDB (Table S7). We found among the top anti SARS-CoV-2 drug candidates the known reverse transcriptase inhibitor didanosine (Figure 7B, Table S7), which other *in silico* screenings have also suggested as a potential treatment for COVID-19 (Alakwaa, 2020; Cava et al., 2020). Among others, NICEdrug.ch also identified: (1) actodigin, which belongs to the family of cardiotonic molecules proven to be effective against MERS-CoV but without mechanistic knowledge (Ko et al., 2020), (2) three molecules in ginger (6-paradol, 10-gingerol, and 6-shogaol) inhibiting catechol methyltransferase, and (3) brivudine, a DNA polymerase inhibitor that has been used to treat herpes zoster (Wassilew, 2005) and prevent MERS-CoV infection (Park et al., 2019), and NICEdrug.ch suggests it for repurposing (Figure S5, Table S7).

**Figure 7.**
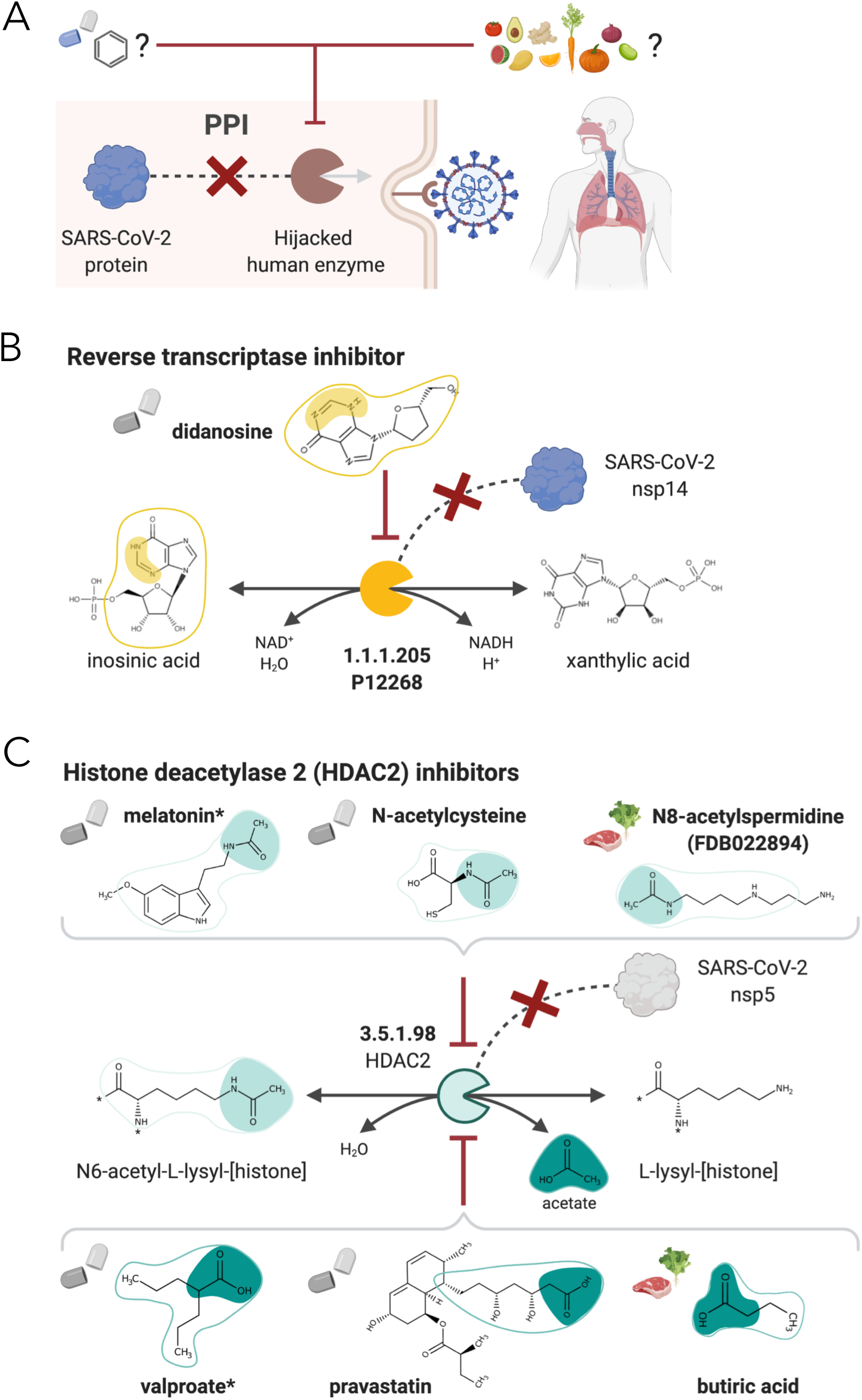
NICEdrug strategy to fight COVID-19, and NICEdrug candidate inhibitors of SARS-CoV-2 host factors: reverse transcriptase and HDAC2. **(A)** Schema of NICEdrug strategy to target COVID-19, wherein a drug (top-left) or molecules in food (top-right) efficiently inhibit a human enzyme hijacked by SARS-CoV-2. Inhibition of this host factor reduces or abolishes protein-protein interactions (PPI) with a viral protein and prevents SARS-CoV-2 proliferation. **(B)** Inhibition of the reverse transcriptase (E.C: 1.1.1.205 or P12268) and the PPI with SARS-CoV-nsp14 by didanosine based on NICEdrug.ch. **(C)** Inhibition of the HDAC2 (E.C: 3.5.1.98) and the PPI with SARS-CoV-nsp5 by molecules containing acetyl moiety (like melatonin, N-acetylcysteine, and N8-acetylspermidine), and molecules containing carboxylate moiety (like valproate, stains, and butyrate) based on NICEdrug.ch. See also Figure S5, Figure S6, Table S7, and Table S8.

Drugs like remdesivir, EIDD-2801, favipiravir, and inhibitors of angiotensin converting enzyme 2 (ACE2) have been used to treat COVID-19 (Jeon et al., 2020), and act through a presumably effective inhibitory mechanism (Figure S6A). For instance, the three drugs remdesivir, EIDD-2801, and favipiravir are believed to inhibit the DNA-directed RNA polymerase (E.C: 2.7.7.6). Here, we used the NICEdrug reactive site-centric fingerprint to seek for alternative small molecules in NICEdrug.ch and fooDB that could be repurposed to target ACE2 and DNA-directed RNA polymerase. NICEdrug.ch identified a total of 215 possible competitive inhibitors of ACE2. Among those is captopril, a known ACE2 inhibitor (Kim et al., 2003), and D-leucyl-N-(4-carbamimidoylbenzyl)-L-prolinamide, a NICEdrug.ch suggestion for drug repurposing to treat COVID-19. We also found 39 food-based molecules with indole-3-acetyl-proline (a molecule in soybean) as top ACE2 inhibitor candidate (Figure S6A, Table S8). To target the same enzyme as remdesivir, EIDD-2801, and favipiravir, NICEdrug.ch identified 1115 inhibitors of the DNA-directed RNA polymerase, like the drug vidarabine, which shows broad spectrum activity against DNA viruses in cell cultures and significant antiviral activity against infections like the herpes viruses, the vaccinia virus, and varicella zoster virus (Suzuki et al., 2006). We further found 556 molecules in food that might inhibit DNA-directed RNA polymerase, like trans-zeatin riboside triphosphate (FDB031217) (Table S8).

One of the host factors identified by Gordon and co-workers is the histone deacetylase 2 (HDAC2) (Gordon et al., 2020), which acetylates proteins and is an important transcriptional and epigenetic regulator. The acetyl and carboxylate moieties are the reactive sites of the forward (N6-acetyl-L-lysyl-[histone]) and reverse (acetate) biotransformation of HDAC2, respectively (Figure 7). NICEdrug.ch recognized a total of 640 drugs for repurposing that can inhibit HDAC2, including 311 drugs sharing the acetyl moiety and showing a NICEdrug score above 0.5 with respect to N6-acetyl-L-lysyl-[histone], and 329 drugs sharing the carboxylate moiety and presenting a NICEdrug score above 0.5 with acetate (STAR Methods). Among the drugs sharing the acetyl reactive site, we identified the known HDAC2 inhibitor melatonin (Wu et al., 2018), and to-our-knowledge new candidates like N-acetylhistamine and N-acetylcysteine. We also located 22 molecules in food with potential HDAC2 inhibitory activity, like N8-acetylspermidine (FDB022894) (Figure 7C, Table S8). Drugs sharing the carboxylate reactive site (as identified with NICEdrug) include the known HDAC2 inhibitors valproate, butyrate, phenyl butyrate (Abdel-Atty et al., 2014) and statins (Kong et al., 2004) (Figure 7C, Table S8). Interestingly, statins have been shown to have protective activity against SARS-CoV-2 (Lodigiani et al., 2020; Zhang et al., 2020). In addition and excitingly, the NICEdrug.ch candidate N-acetylcysteine is a commonly used mucolytic drug that is sometimes considered as a dietary supplement and has putative antioxidant properties. Indeed, N-acetylcysteine is believed for long to be precursor of the cellular antioxidant glutathione (Mårtensson et al., 1989), but has unknown pharmacological action. NICEdrug.ch suggests that N-acetylcysteine might present a dual antiviral activity: firstly, N-acetylcysteine is converted to cysteine by HDAC2 and by that means, it is competitively inhibiting the native function of HDAC2 and interactions with viral proteins (Figure 7C, Table S8). Cysteine next fuels the glutathione biosynthesis pathway and produces glutathione in two steps.

Given the high coverage of validated molecules with activity against SARS-CoV-2 that NICEdrug.ch captured in this unbiased and reactive site-centric analysis, we suggest there might be other molecules in the set of 1,300 NICEdrug.ch candidates that could also fight COVID-19. Excitingly, there are many molecules that can be directly tested since these are drugs that have already passed all safety regulations or are molecules in food, like N-acetylcysteine for which we further reveal an action mechanism behind its potential anti SARS-CoV-2 activity. Other new candidates for which no safety data is available should be further validated experimentally and clinically. The mechanistic analyses provided by NICEdrug.ch could also guide new pharmacokinetic and pharmacodynamic models simulating SARS-CoV-2 infection and treatment.

## Discussion

To systematically illuminate the metabolism and all enzymatic targets (competitively inhibited) of known drugs and hypothetical prodrugs to aid in the development of new therapeutic compounds, we used a proven reaction-prediction tool BNICE.ch (Hatzimanikatis et al., 2005) and an analysis of neighboring atoms of reactive sites analogous to BridgIT (Hadadi et al., 2019) and performed the first large-scale computational analysis of drug biochemistry and toxicity in the context of human metabolism. The analysis involved over 250,000 small molecules, and curation and computation of bio- and physicochemical drug properties that we assembled in an open-source drug database NICEdrug.ch that can generate detailed drug metabolic reports and can be easily accessed and used by researchers, clinicians, and industry partners. Excitingly, NICEdrug.ch revealed 20 million potential reactive sites at the 250,000 small molecules of the database, and there exist over 3,000 enzymes in the human metabolism that can be inhibited with the 250,000 molecules. This is because NICEdrug.ch can identify *all* potential metabolic intermediates of a drug and scans these molecules for substructures that can interact with catalytic sites across all enzymes in a desired cell.

NICEdrug.ch adapts the metric previously developed for reactions in BridgIT (Hadadi et al., 2019) to precisely compare drug-drug and drug-metabolite pairs based on similarity of reactive site and the neighborhood around this reactive site, which we have recently shown outperforms previously defined molecular comparison metrics (Hadadi et al., 2019). Since NICEdrug.ch shows high specificity in the identification of such reactive sites and neighborhood, it provides a better mechanistic understanding than currently available methods (Robertson, 2005). Despite these advances, it remains challenging to systematically identify non-competitive inhibition or targeting of non-enzymatic biological processes. We suggest coupling NICEdrug.ch drug metabolic reports with other *in silico* and experimental analyses accounting for signaling induction of small molecules and other non-enzymatic biological processes like transport of metabolites in a cell. The combined analysis of drug effects on different possible biological targets (not uniquely enzymes) will ultimately increase the coverage of molecules for which a mechanistic understanding of their mode of action is assigned.

A better understanding of the mechanisms of interactions and the specific nodes where the compounds act can help re-evaluate pharmacokinetic and pharmacodynamic models, dosage, and treatment. Such understanding can be used in the future to build models that correlate the pharmacodynamic information with specific compounds and chemical substructures in a manner similar to the one used for correlating compound structures with transcriptomic responses. We have shown for one of the most commonly used anticancer drugs, 5-FU, that NICEdrug.ch identifies and ranks alternative sources of toxicity and hence can guide the design of updated models and treatments to alleviate the drug’s side-effects.

The mechanistic understanding will also further promote the development of drugs for repurposing. While current efforts in repurposing capitalize on the accepted status of known drugs, some of the issues with side effects and unknown interactions limit their development as drugs for new diseases. Given that drug repurposing will require new dosage and administration protocols, the understanding of their interactions with the human metabolism will be very important in identifying, developing, and interpreting unanticipated side effects and physiological responses. We evaluated the possibility of drug repurposing with NICEdrug.ch as a substitute for statins, which are broadly used to reduce cholesterol but have many side effects. NICEdrug.ch and its reactive site-centric comparison accurately cluster both family types of statins, even though they are similar in overall molecular structure and show different reactivity. In addition, NICEdrug.ch suggests a set of new molecules with hypothetically less side effects (Endo and Hasumi, 1993; Tanaka et al., 2004) that share reactive sites with statins.

A better mechanistic understanding of drug targets can guide the design of treatments against infectious diseases, for which we need effective drugs that target pathogens without side effects in the host cell. This is arguably the most challenging type of problem in drug design, and indeed machine learning has continuously failed to guide such designs given the difficulty in quantifying side effects—not to mention in acquiring large, consistent, and high-quality data sets from human patients. To demonstrate the power of NICEdrug.ch for tackling this problem, we sought to identify drugs that target liver-stage malaria parasites and minimize the impact on the human host cell. We identified over 500 drugs that inhibit essential *Plasmodium* enzymes in the liver stages and minimize the impact on the human host cell. Our top drug candidate is shikimate 3-phosphate targeting the parasite’s shikimate metabolism, which we recently identified as essential in a high-throughput gene knockout screening in *Plasmodium* (Stanway et al., 2019). Excitingly, our suggested antimalarial candidate shikimate 3-phosphate has already been used for *Escherichia* and *Streptococcus* infections without appreciable side effects (Díaz-Quiroz et al., 2018).

Finally, minimizing side effects becomes especially challenging in the treatment of viral infections, since viruses fully rely on the host cell to replicate. As a last demonstration of the potential of NICEdrug.ch, we sought to target COVID-19 by identifying inhibitors of 22 known enzymatic host factors of SARS-CoV-2 (Gordon et al., 2020). NICEdrug.ch identified over 1,300 molecules that might target the 22 host factors and prevent SARS-CoV-2 replication. As a validation, NICEdrug.ch correctly identified known inhibitors of those enzymes, and further suggested safe drugs for repurposing and other food molecules with activity against SARS-CoV-2. Among the NICEdrug.ch suggestions for COVID-19, based on the knowledge on its mechanism and safety, we highlight N-acetylcysteine as an inhibitor of HDAC2 and SARS-CoV-2.

Overall, we believe that a systems level or metabolic network analysis coupled with an investigation of reactive sites will likely accelerate the discovery of new drugs and provide additional understanding regarding metabolic fate, action mechanisms, and side effects and can complement on-going experimental effects to understand drug metabolism (Javdan et al., 2020). We suggest the generation of drug metabolic reports to understand the reactivity of new small molecules, the possibility of drug repurposing, and the druggability of enzymes. Our results using NICEdrug.ch suggest that this database can be a novel avenue towards the systematic pre-screening and identification of drugs and antimicrobials. In addition to human metabolic information, NICEdrug.ch currently includes information for the metabolism of *P. berghei* and *E. coli*. Because we are making it publicly available (https://lcsb-databases.epfl.ch/pathways/Nicedrug/), our hope is that scientists and medical practitioners alike can make use of this unique database to better inform their research and clinical decisions—saving time, money, and ultimately lives.

## Acknowledgements

We would like to thank Dr. Ljubisa Miskovic, Dr. Volker T. Heussler, Dr. Reto Caldelari, Dr. Jens Nielsen, and Dr. Adil Mardinoglu for critical feedback and discussions. The graphical abstract and Figures 6, 7, S5, and S6 were done with BioRender. H.M. was funded by the European Union’s Horizon 2020 research and innovation program under the Marie Sklodowska-Curie grant agreement No 72228. H.M., K.H. and J.H. were supported by the Ecole Polytechnique Fédérale de Lausanne (EPFL). A.C. and V.H. were funded by grant 2013/155 (MalarX), N.H. and J.H. were funded by grant 2013/158 (MicroScapesX); both are SystemsX.ch grants awarded by the SNSF.

## Author Contributions

Conceptualization, V.H., H.M., N.H., A.C-P.; Methodology, V.H., H.M., K.H., N.H., A.C-P.; Software, H.M., K.H., J.H.; Formal analysis, H.M., K.H.; Investigation, H.M., K.H., A.C-P.; Writing-Original Draft, H.M., A.C-P.; Writing-Review & Editing, V.H., H.M., A.C-P., J.H., N.H.; Visualization, H.M., A.C-P.; Supervision, V.H.; Project Administration, V.H., H.M.; Funding Acquisition, V.H.

## Supplementary figure title and legends

**Supplementary figure 1.**
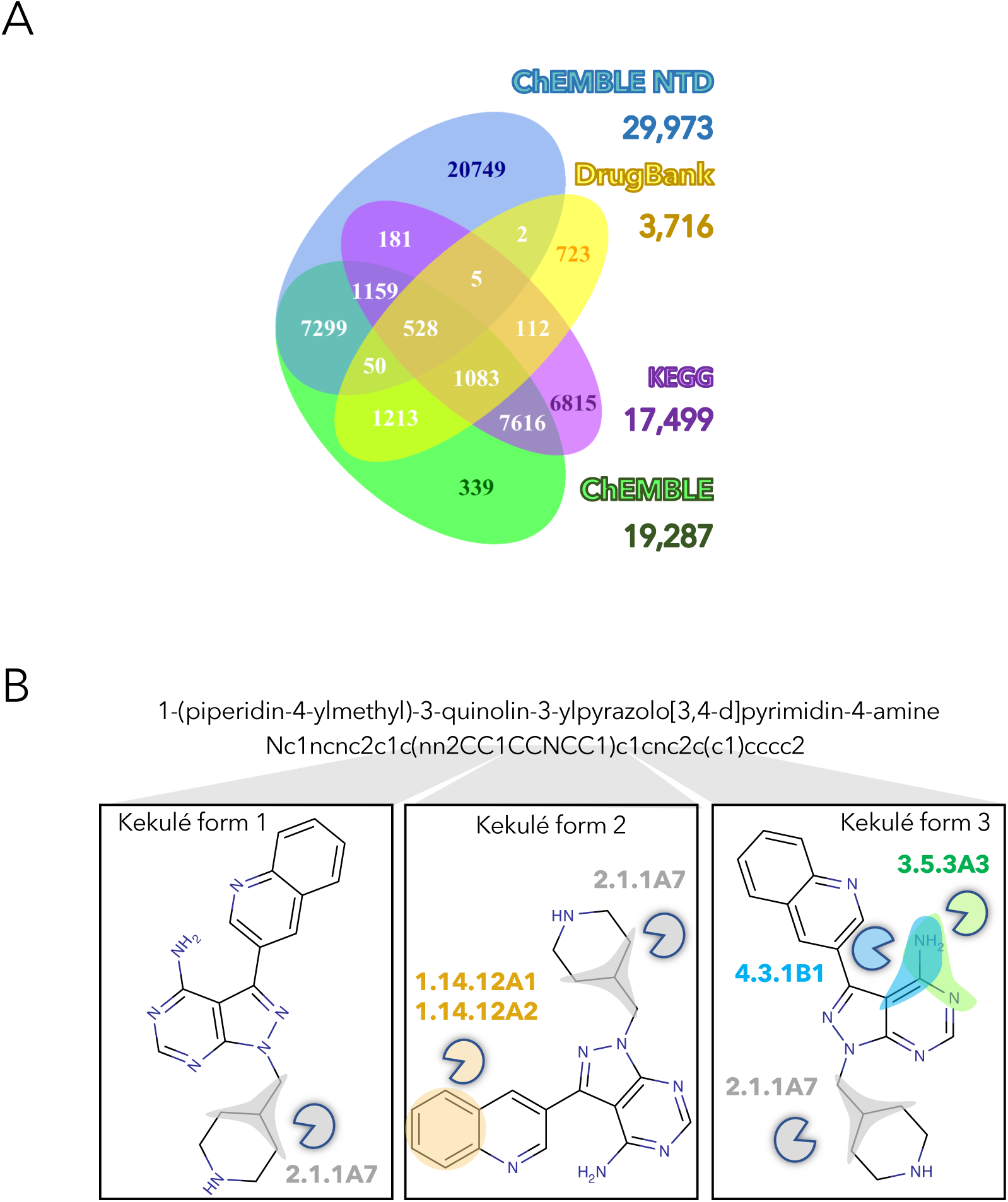
Overview of number of molecules in NICEdrug.ch and their structural curation, related to Figure 1. **(A)** Venn diagram showing the number of compounds in NICEdrug.ch and their source database: KEGG, DrugBank, ChEMBLE NTD, and ChEMBLE. **(B)** Representation on how different kekulé forms affect the identification of reactive sites and prediction of biological activity for an example molecule.

**Supplementary figure 2.**
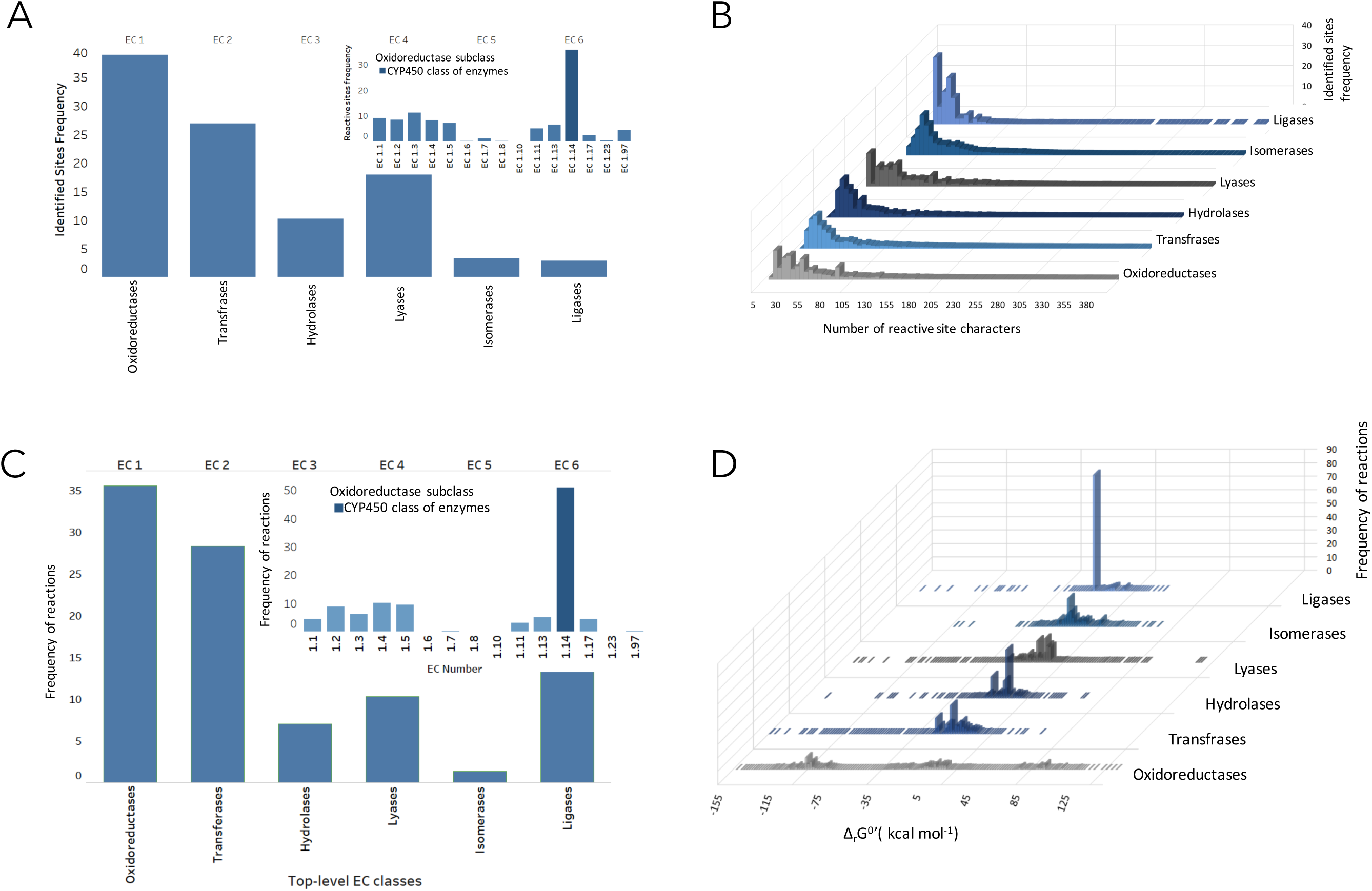
Distribution of reactive sites and metabolic reactions as of E.C. numbers linked to all molecules in NICEdrug.ch, related to Figure 1. **(A)** Distribution of reactive sites identified in all molecules of NICEdrug.ch among classes of E.C. numbers. **(B)** Specificity of reactive sites identified in drugs based on length and types of participating atoms. **(C)** Distribution of drug metabolic reactions based on class of E.C. number. **(D)** Distribution of Gibbs free energy for the drug metabolic reactions, which are the reactions linked to all molecules of NICEdrug.ch.

**Supplementary figure 3.**
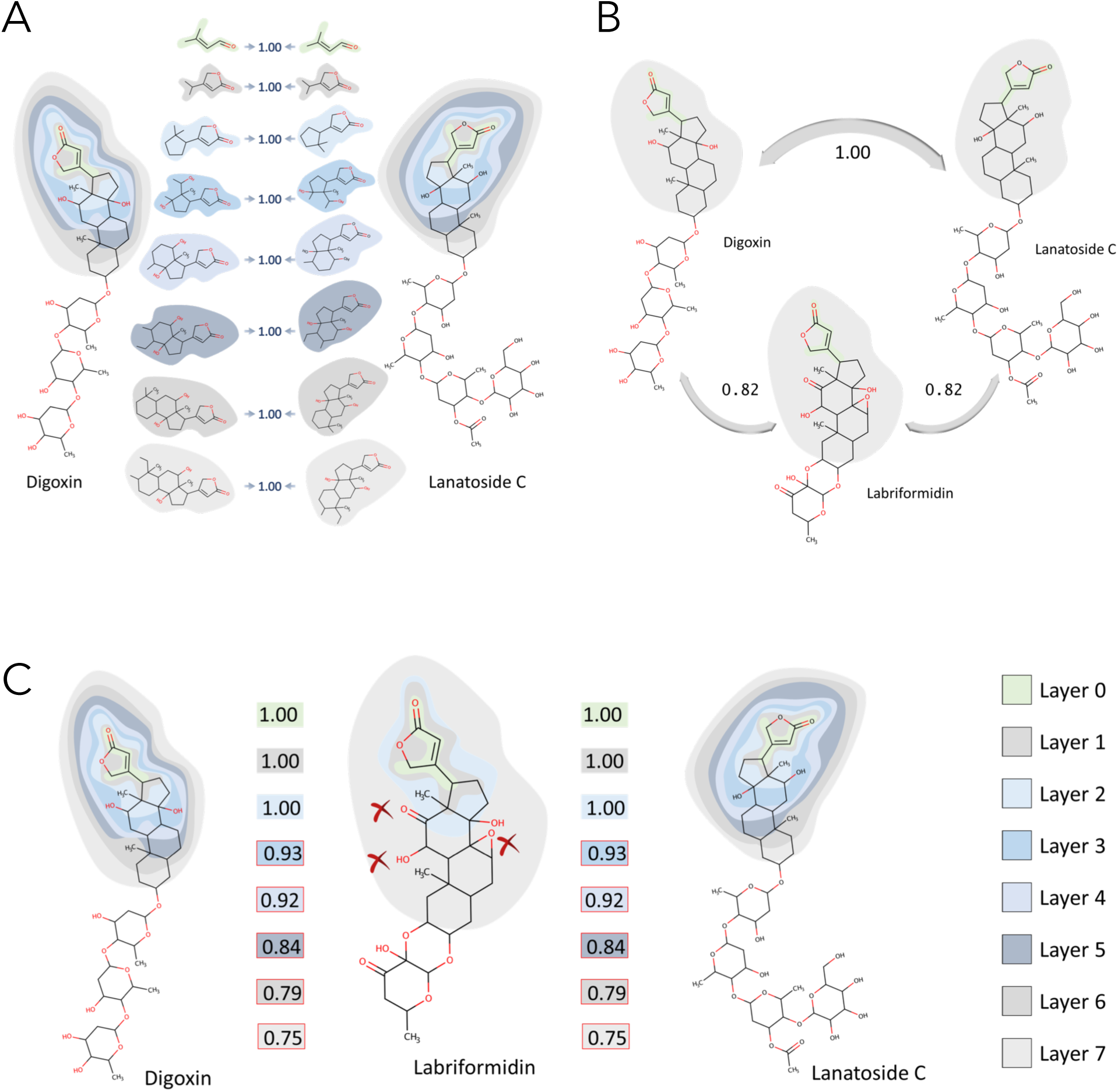
Description of NICEdrug score, related to Figures 1, 2, 3, 4, 5, and 6. Example of NICEdrug score calculation. The NICEdrug score takes into account the structure of a molecule’s reactive site and its seven-atom-away neighborhood for similarity evaluation, analogous to BridgIT.

**Supplementary figure 4.**
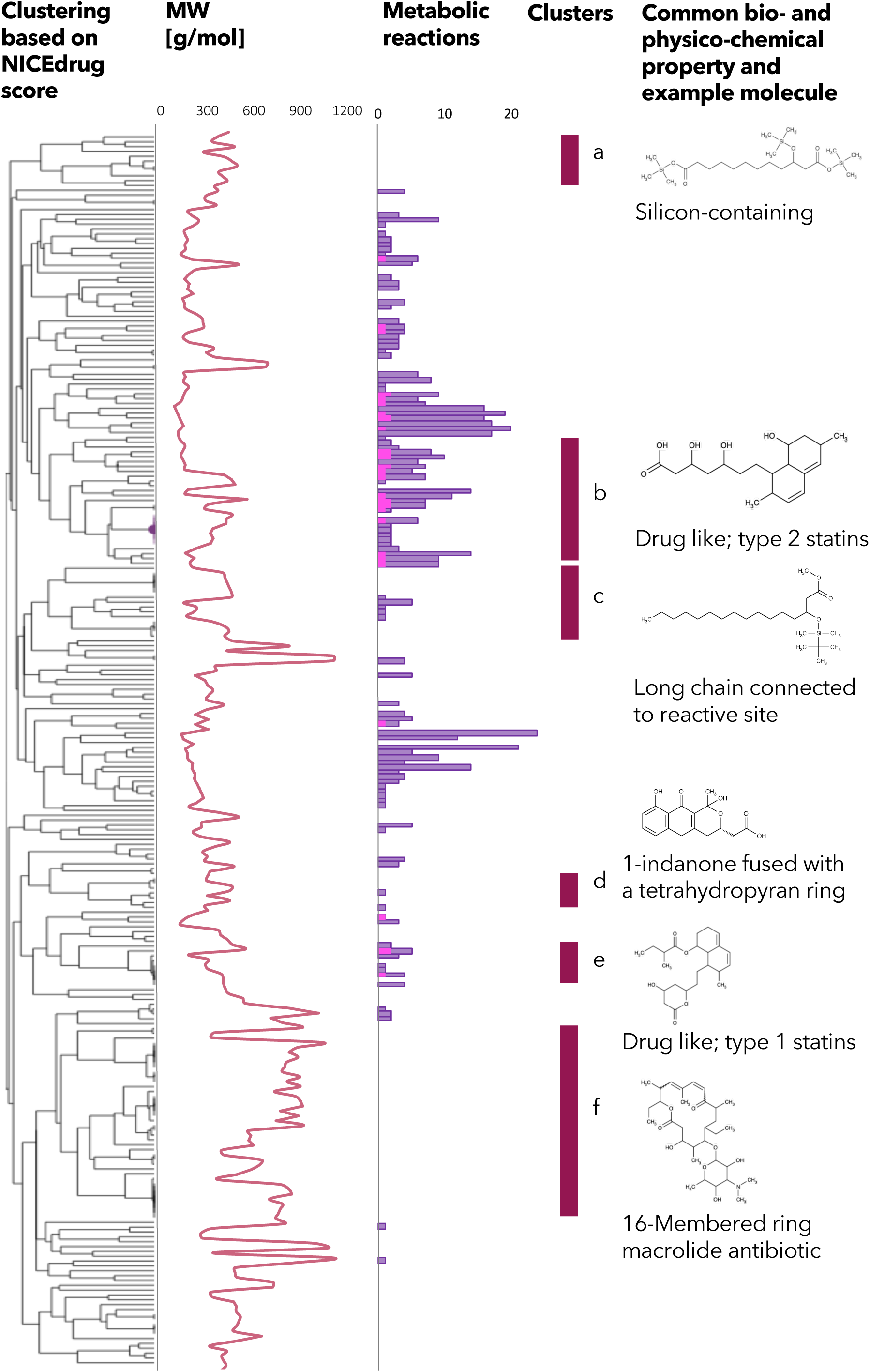
Clustering based on NICEdrug score, molecular weight, and reactivity of statin like molecules, related to Figure 5. Hierarchical clustering based on the NICEdrug score of all molecules in NICEdrug.ch that contain statin reactive site (left). We report the molecules’ molecular weight (middle left) and number of drug metabolic reactions or reactions in which these drugs participate (middle). The molecular weight seems to be inversely correlated with the number of drug metabolic reactions. We highlight six clusters of drugs (a-f, middle right) and an example representative molecule (left). Interestingly, these clusters also group molecules based on bio- or physico-chemical properties: “cluster a” involves a range of silicon-containing chemical molecules, “cluster b” are drug like molecules of type 2 statins, “cluster c” includes chemical molecules with a long chain connected to the reactive site, “cluster d” involves molecules with 1-indanone fused with a tetrahydropyran ring, “cluster e” comprises drug-like molecules of type 1 statins, and “cluster f” are 16-membered ring macrolide antibiotics.

**Supplementary figure 5.**
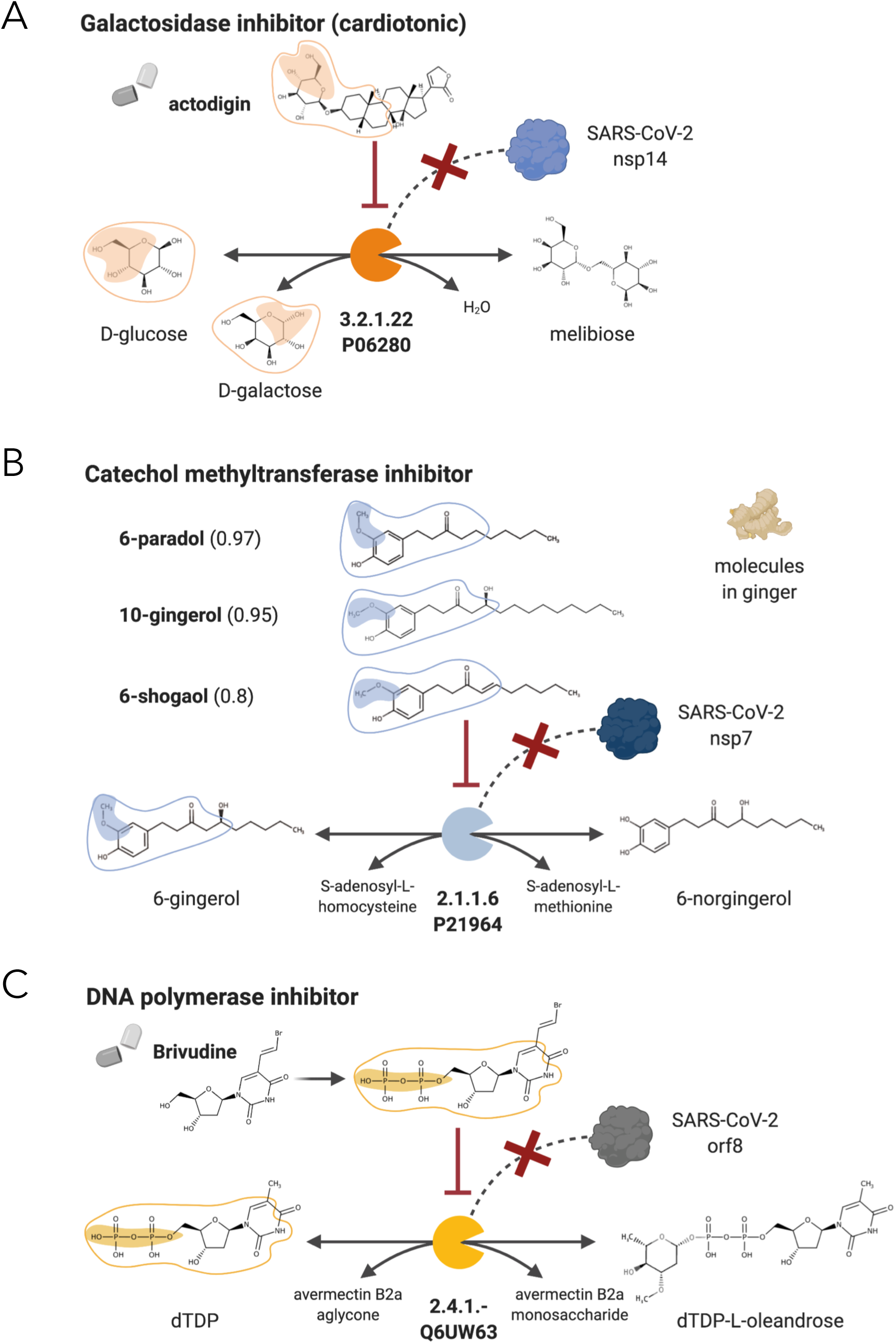
NICEdrug candidate inhibitors of SARS-CoV-2 host factors: galactosidase, catechol methyltransferase, and DNA polymerase, related to Figure 7. **(A)** Inhibition of the galactosidase (E.C: 3.2.1.22 or P06280) and the PPI with SARS-CoV-2 nsp14 by actodigin based on NICEdrug.ch. **(B)** Inhibition of the catechol methyltransferase (E.C: 2.1.1.6 or P21964) and the PPI with SARS-CoV-2 nsp7 by 6-paradol, 10-gingerol, and 6-shogaol, which are molecules in ginger, based on NICEdrug.ch. **(C)** Inhibition of the DNA polymerase (E.C: 2.4.1.-) and the PPI with SARS-CoV-2 nsp8 by brivudine based on NICEdrug.ch.

**Supplementary figure 6.**
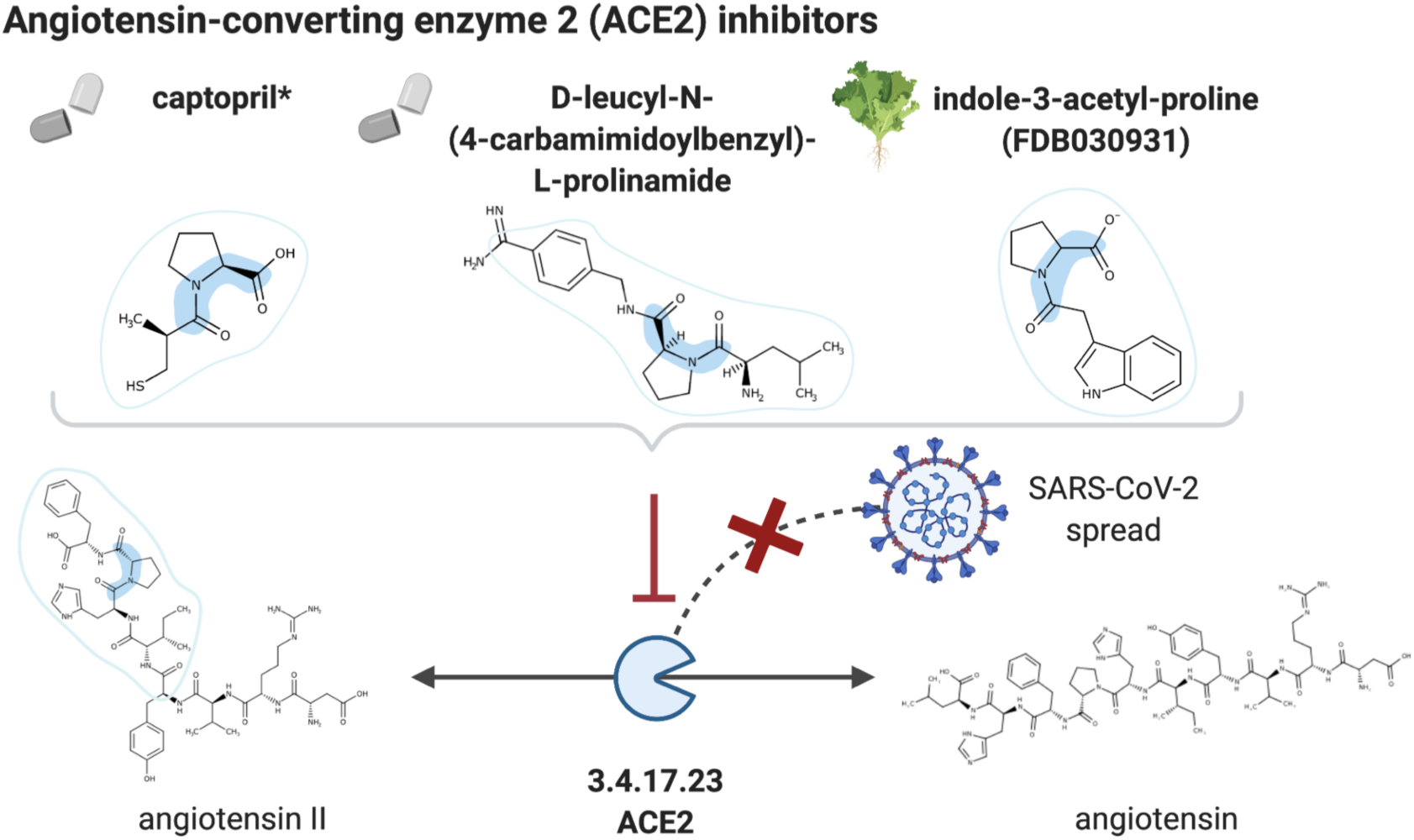
NICEdrug candidate inhibitors of ACE2, related to Figure 7. Inhibition of the ACE2 (E.C: 3.4.17.23), a putative host factor of SARS-CoV-2, by the known inhibitor captopril, and NICEdrug candidates D-leucyl-N-(4-carbamimidoylbezyl)-L-prolinamide and indole-3-acetyl-proline.

## Materials and Methods

### Representation of metabolic neighborhood in figures of this manuscript

We represent the metabolic neighborhood of a drug with reactions or steps away (arrows), where each step away (circle connected to arrow) involves a set of compounds. We extract compounds at each step that present a high NICEdrug score (value under metabolite name) with the native substrate of a reaction in the human cell. Reactive sites common to neighbor metabolites and native human metabolites are shaded with colors matching the color of the enzymes (packmen) that are inhibited. The neighborhood (seven atoms away, as considered in NICEdrug score) of the reactive sites is circled in the metabolites and native human metabolites with the same color as the reactive sites and enzymes. Compounds marked with * are confirmed inhibitors and references are provided in the main text.

### Representation of enzymatic inhibition in figures of this manuscript

We represent the enzymes and catalyzed reactions inhibited by NICEdrug candidates. Highlighted are the reactive site and neighborhood (as considered in the NICEdrug score) in candidate drugs and metabolites, which are native substrates of the human enzymes. The SARS-CoV-2 proteins interaction with the enzyme is presumed to be diminished or abolished upon inhibition of the human enzyme. Compounds marked with * are confirmed inhibitors and references are provided in the main text.

### Curation of input molecules used in the construction of NICEdrug.ch

We constructed the NICEdrug.ch database to gather small molecules suitable for treatment of human diseases. We collected the SMILES structure, synonyms, and any available bio- and physico-chemical property included from three source databases: KEGG, ChEMBL, and DrugBank, which added up to 70,976 molecules by January 2018 (Figure S1A). Only molecules that were fully structured were imported to our database. We further curated the imported molecules by removing duplicate structures and merging annotations from different databases into one molecule entry in the database. For removing duplicate structures we used canonical SMIELS (Weininger, 1988) generated by openbabel (O’Boyle et al., 2011) version 2.4.0. This unification method is based on atoms and their connectivity in a molecule in terms of a molecular graph that is captured by the canonical SMILES. Therefore, different resonance forms, stereoisomers, as well as dissociated and charged states of the same compound are mapped to one entry in database. Furthermore, we filtered molecules based on Lipinski rules (Lipinski et al., 2001): (1) the molecular weight should be less than 500 Dalton, (2) the number of hydrogen bond donors should be less than five, (3) the number of hydrogen bond acceptors should be less than ten, and (4) an octanol-water partition coefficient (log P) should be less than five. According to Lipinski rules an active orally drug does not violate more than one of the above criteria. We calculated criteria one, two and three based on the structural information from SMILES of molecules. To assess criterion four, we relied on reported data in the source database. We kept in the NICEdrug.ch database those compounds for which the partition coefficient was not available.

We performed a separate analysis to account for non-unique graph representations of aromatic rings, also called *kekulé structures*. The existence of aromatic rings and the fact that bond-electrons are shared within the ring make several single-double bond assignments possible, which results in multiple kekulé representations for a single molecule (Figure S1B). We included all such kekulé structures to account for alternative atom-bond connectivity and associated reactivity. We call “effective forms” to the kekulé representations that show different reactive sites than their canonical structures. For example, there can be two effective forms plus the canonical structure (Figure S1B). In total, we found 42,092 effective forms for 29,994 aromatic compounds in NICEdrug.ch database and we kept them for further analysis.

We also computed the thermodynamic properties of all drugs in NICEdruch.ch. Specifically, we computed the Gibbs free energy of formation (Δ_*f*_*G*′°) using the group contribution method of Mavrovouniotis (Jankowski et al., 2008).

The NICEdrug.ch database includes a total number of 48,544 unique and curated small molecules (Figure S1A).

### Identification of reactive sites in drugs

The 3D structures of enzyme pockets are complex and mostly unknown. Therefore, evaluating and comparing docking of two small molecules in the pocket of a specific target is impossible most of the times. Using BNICE.ch, we focused on the complementary structure of active sites on substrates, also called *reactive site*. To recognize the potential reactive sites on molecules, we scanned molecules using expert-curated generalized reaction rules of BNCIE.ch (Hadadi et al., 2016), which mimic the identification of substrates by the enzyme pocket and account for the promiscuous activity of enzymes. Theses reaction rules incorporate the information of biochemical reactions and have third-level Enzyme Commission (EC) identifiers. Each BNICE.ch reaction rule accounts for three levels of information: (1) atoms in reactive sites of compounds, (2) connectivity and configuration of atom bonds in the reactive site, and (3) mechanism of bond breakage and formation during the reaction. As of May 2020, BNICE.ch contains 450 bidirectional generalized reaction rules that can reconstruct 8118 KEGG reactions (Hadadi et al., 2016). Here, we include all BNICE.ch rules to identify all possible reactive sites on a given drug in two steps. First, a BNICE.ch rule identifies all atoms in a compound that belong to the rule’s reactive site. Second, the rule evaluates the connectivity of the atoms previously identified. The candidate compounds for which a BNICE.ch rule identified a reactive site were validated as metabolically reactive and considered for analysis in NICEdrug.ch.

It is important to note that thanks to the generalized reaction rules, which abstract the knowledge of thousands of biochemical reactions, BNICE.ch is able to reconstruct known biotransformations and also propose novel metabolic reactions. This was demonstrated in the reconstruction of the ATLAS of Biochemistry (Hadadi et al., 2016), which involves up to 130,000 reactions between known compounds.

### Analysis of drug metabolism in human cells

To mimic biochemistry of human cells and simulate human drug metabolism, we collected all available information (metabolites and metabolic activities or EC numbers of enzymes) on human metabolism from three available databases: the human metabolic models Recon3D (Brunk et al., 2018) and HMR (Pornputtapong et al., 2015), and the Reactome database (Croft et al., 2011). These three databases include a total of 2,266 unique human metabolites and 2,066 unique EC numbers of enzymes (Table S1).

To explore the biochemical space beyond the known human metabolic reactions and compounds, we used (1) the generalized enzymatic reaction rules of BNICE.ch that match up to the third EC level the collected human enzymes, and (2) all of the collected human metabolome. We evaluated the reactivity of each drug in a human cell using the retro-biosynthesis algorithm of BNICE.ch, which predicts hypothetical biochemical transformations or *metabolic neighborhood* around the drug of study. We generated with BNICE.ch metabolic reactions in which each drug and all known human metabolites could participate as substrate or products. We also allowed a set of 53 known cofactors to react with the human metabolites (Table S1).

We define the boundaries of the metabolic neighborhood of a molecule with a maximum number of reactions or *steps away* that separate the input molecule (drug of study) from the furthest compound. In BNICE.ch, a generation *n* of compounds involves all metabolites that appear for the first time in the metabolic neighborhood of a drug after *n* reactions or steps happened. For example, in the case study of 5-FU we find the compound 5-Fluorouridine in generation 2 or 2 steps away, which means there are two metabolic reactions that separate 5-FU and 5-Fluorouridine (Figure 2).

In NICEdrug.ch, there exist 197,246 compounds in generation 1 (1 step away) from all input drugs. The 197,246 compounds are part of the potential drug metabolic neighborhood in human cells. Out of all generation 1 molecules, 13,408 metabolites can be found in human metabolic models and HMDB database (Wishart et al., 2018), 16,563 metabolites exist in other biological databases, and the remaining 167,245 metabolites are catalogued as known compounds in chemical databases (i.e., PubChem). Note that HMDB includes native human metabolites and non-native human compounds, like food ingredients.

The 197,246 products that are one-step away of all NICEdrug.ch molecules are part of a hypothetical biochemical neighborhood of 630,449 drug metabolic reactions. Of all drug metabolic reactions, 5,306 reactions are cataloged in biological databases, and the remaining 625,143 reactions are novel. A majority of the reactions involved oxidoreductases (42.54%), broken down into 27.45% of lyases, 7.15% of hydrolases, 6.28% of transferases, 1% of isomerases, and 15.58% of ligases. Interestingly based on the previously identified reactive sites, out of the 265,935 (42.54% of 625,143) oxidoreductase reactions, 49.92% are catalyzed by the p450 family of enzymes, which are known to be responsible for the metabolism of drug (Figure S2C).

### Using NICEdrug.ch database for analysis of the metabolic neighborhood of a drug

In NICEdrug.ch webserver, users can look up for a drug using the drugs’ name and other identifiers like ChEMBL, DrugBank and KEGG. NICEdrug.ch will report a unique identifier for the compound that will be input for upcoming analysis modules. The *predict metabolism* module allows to study the metabolic network around an input molecule. The input to this module is: (1) the unique identifier of the drug of interest, (2) a maximum number of reactions or *steps away* that shall separate the input drug to the furthest compound in the metabolic neighborhood.

The output of this analysis is a report in the form of a csv file that includes all compounds and metabolic reactions in the metabolic neighborhood of the input drug. One can also export the neighborhood in the form of a visual graph, in which nodes are molecules and edges are reactions.

### Definition of the NICEdrug score

Based on the theory of lock and key, two metabolites that can be catalyzed by the same enzyme may have similar reactive sites and also neighboring atoms. In order to quantify the similarity inside and around reactive sites of <u>two molecules</u>, we developed a metric called *NICEdrug score* (Figure S3), which is inspired on BridgIT (Hadadi et al., 2019). BridgIT assesses the similarity of <u>two reactions</u>, considering the reactive site of the participating substrates and their surrounding structure until the seventh atom out of the reactive site.

The NICEdrug score is an average of two similarity evaluations: (1) the atom-bond configuration inside reactive site (α parameter), and (2) the 7 atom-bond chain molecular structure around the reactive site (β parameter). The NICEdrug score, and its parameters α and β, range between 0 and 1 when they indicate no similarity and identical structure, respectively. Different constraints on the α and β parameters determine the identification of different types of inhibition like para-metabolites and anti-metabolites (see other sections in this Materials and Methods).

We show the evaluation of NICEdrug scores for three example compounds (Figure S3). In this example, Digoxin, Labriformidin and Lanatoside C all share the reactive site corresponding to EC number 5.3.3.- (α=1). Starting from the atoms of the identified reactive site, eight description layers of the molecule were formed, where each layer contains a set of connected atom-bond chains. Layer zero includes types of atoms of reactive site and their count. Layer 1 expands one bond away from all of the atoms of reactive site and accounts for atom-bond-atom connections. This procedure is continued until layer 7, which includes the sequence of 8 atoms connected by 7 bonds. Then, we compare the fingerprint of each molecule to the other participants of the class based on the Tanimoto similarity scores. A Tanimoto score near 0 designates no or low similarity, whereas a score near 1 designates high similarity in and around reactive site. Lanatoside C and Digoxin share the same substructure till 8 layers out of reactive site which is presented in the NICEdrug score by preserving score 1 in all layers, so the overall Tanimoto score for these two compounds in the context of EC number 5.3.3.-is 1 (α=1 and β=1). However, the structure of two compounds are not exactly the same and actually Lanatoside C has 8 more carbon atoms and 6 more oxygen atoms, shaped as an extra benzenehexol ring and an ester group. Although this part is far from the reactive site, based on the NICEdrug score they both can perfectly fit inside the binding pocket of a common protein related to this reactive site. This hypothesis is proved by experiments reported in KEGG and DrugBank. According to DrugBank and KEGG, Lanatoside C has actions similar to Dioxin and both of them have the same target pathways: Cardiac muscle contraction and Adrenergic signaling in cardiomyocytes. Furthermore, target protein for both of them is ATP1A.

Also, the NICEdrug score effectively captures and quantifies differences around the reactive site. The substructure around the reactive site in Labriformidin is slightly different (α=1 and β <1). The difference is calculated trough different layers of the NICEdrug score.

In the case study of 5-FU, in order to predict competitive inhibition, we analyzed all the metabolites that share reactive site with 5-FU or its downstream products (α=1) and then we ranked the most similar metabolites based on their similarity in neighborhood of reactive site to 5-FU or its downstream products (β). To assess the structural differences in the reactive sites themselves (α), we implemented the Levenshtein edit distance algorithm (Levenshtein, 1966) to determine how many deletions, insertions, or substitutions of atom/bonds are required to transform one pattern of reactive site into the other one. Here, the edit distance explains the difference between the reactive sites of the intermediate and the human metabolite. However, even slight changes in the reactive site affect its interaction with the binding site. To ensure that the divergence retained the appropriate topology, we compared the required edit on reactive site with interchangeable groups, termed bioisosteric groups (Papadatos and Brown, 2013). These bioisosteric groups contain similar physical or chemical properties to the original group and largely maintain the biological activity of the original molecule. An example of this is the replacement of a hydrogen atom with fluorine, which is a similar size that does not affect the overall topology of the molecule. For this analysis, we used 12 bioisosteric groups adapted from the study by Papadatos et.al. (Papadatos and Brown, 2013).

To predict irreversible Inhibitors in metabolism of 5-FU, we kept only molecules with a similarity score greater than 0.9 to metabolites (β>0.9), to preserve a high similarity in the neighborhood of the reactive sites. Then, we checked which ones contained reactive sites that differed only in the replacement of bioisosteric groups (α∼1).

### Classification of drugs based on the NICEdrug score

Classification of compounds with similar structure is normally used to assign unknown properties to new compounds. For instance, one can infer ligand-protein binding for a drug when its action mechanism or the structure of the target proteins are not known. In this study, we have demonstrated four strategies to classify drugs (Figure 1), which are from less to more stringent: classifying (1) molecules that participate in reactions with the same EC up to the 3^th^ level, (2) molecules that in addition share a BNICE.ch reaction rule, (3) molecules that in addition to both previous points share reactive site, (4) molecules that show high similarity of reactive site and neighborhood based on the NICEdrug score.

The EC number guarantees that molecules are catalyzed with similar overall reaction mechanism. Generalized reaction rules from BNICE.ch further capture different submechanisms inside an EC number (Hadadi et al., 2016). A BNICE.ch reaction rule might involve more than one reactive site. Hence, information of reactive sites provide further insights into the molecule’s reactivity. Furthermore, similarity of reactive sites and their neighborhoods based on the NICEdrug score increase the comparison resolution and this is the basis of the classification in NICEdrug.ch.

In NICEdrug.ch database there exist 95,342 classes that comprise all drugs and human compounds sharing EC, BNICE.ch rule, and reactive site (classification based on our strategy 3). We computed the NICEdrug score between all pairs of molecules in a class and this information is available in NICEdrug.ch.

### Identification of drugs acting as para-metabolites based on NICEdrug score

Small molecules that share reactive site and are structurally similar to native human metabolites enter and bind the pocket of native enzymes and competitively inhibiting catalysis acting as para-metabolites (Ariens, 2012). In this study, we define as para-metabolite any drug or any of its metabolic neighbors that (1) shares reactive site with native metabolites (α=1), and (2) preserves a high NICEdrug score with respect to the reactive site neighborhood (β>0.9).

### Identification of drugs acting as anti-metabolites based on NICEdrug score

Small molecules that do not share reactive site but are structurally similar to native human metabolites might enter the binding pocket of native enzymes and inhibiting catalysis acting as anti-metabolites (Ariens, 2012). In this study, we define as anti-metabolite any drug or any of its metabolic neighbors that (1) differs slightly in reactive site from a native metabolite (α∼1), and (2) preserves high similarity in the reactive site neighborhood (β>0.9). We hypothesize that a low divergence in the reactive site, still allows a non-native compound to enter and bind the enzyme pocket since it is structurally similar enough to the native substrate.

### Identification of NICEdrug toxic alerts

We obtained all NICEdrug toxic alters from ToxAlert database (Sushko et al., 2012). ToxAlert database includes about 1,200 structural toxic alerts associated with particular types of toxicity. Toxic alerts are provided in the form of SMART patterns that are searchable in SMILES structure of molecules. NICEdrug.ch uses openbable tool (O’Boyle et al., 2011) to search for these structural alerts on SMILES of compounds.

### Collection of reference toxic molecules in NICEdrug.ch

Studying the adverse effects of chemicals on biological systems has led to development of databases cataloging toxic molecules. The Liver Toxicity Knowledge Base (LTKB) integrates 1,036 molecules annotated with human Drug-induced liver injury risk (severity). Super toxic DB include about 60k toxic molecules, that are annotated with their toxicity estimate, LC_50_/LD_50_ i.e., lethal dose or concentration at which 50% of a population dies.

As a resource of approved toxic molecules, we collected all of the molecules cataloged as toxic in LTKB and super toxic databases. We used this collection as a reference to compare the similarity of drugs or and products of drug metabolism with approved toxic molecules.

### Definition of a toxicity score in NICEdrug.ch

The number of molecules labeled as toxic in databases is disproportionally low compared to the space of compounds. On the other hand, toxic alerts are defined for a big number of compounds and are linked to redundant molecular structures.

We measured the similarity of drugs and their metabolic neighbors with the collection of reference toxic molecules using the NICEdrug score. We assigned toxic alerts to molecules in NICEdrug.ch if a molecule and toxic molecule shared a molecular substructure linked to the toxic alert.

Finally, NICEdrug.ch provides a toxicity report in the form of a csv file for each molecule in the metabolic neighborhood including six values linked to the most similar toxic molecules in both toxic reference databases (LTKB and supertoxic databases): (1) the NICEdrug score between the drug and those most similar toxic molecules, (2) the severity degree of the hepatotoxic compound, and log(LC_50_) of the supertoxic compound, and (3) the number of common toxic alerts between the drug and the most similar toxic molecules. The list of toxic alerts is also provided.

We combined the six values of the toxicity report into a toxicity score defined as follows: 

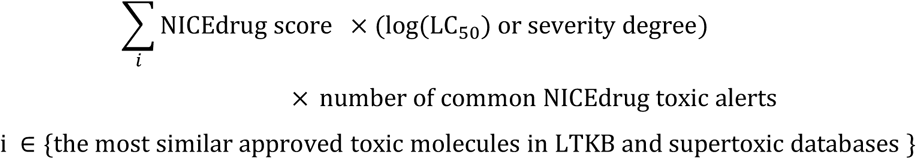

The toxicity score in NICEdrug.ch served to quantify the toxicity of each molecule in the metabolic neighborhood of a drug, recapitulate known toxic molecules, and suggest new toxic compounds (Figure 4).

### Analysis of essential enzymes and linked metabolites in *Plasmodium* and human cells

We extracted information of essential genes and enzymes for liver-stage malaria development from our recent study (Stanway et al., 2019). In this study, we developed the genome-scale metabolic model of *Plasmodium berghei*, which shows high consistency (approximately 80%) with the largest gene knockout datasets in *Plasmodium* blood (Bushell et al., 2017) and liver stages (Stanway et al., 2019). There are 178 essential genes for *P. berghei*’s growth simulating liver-stage conditions (Stanway et al., 2019). Here, we identified the substrates of those essential metabolic enzymes, which comprise a set of 328 metabolites (Table S5). To further minimize on the host cell, we filtered out those *Plasmodium* enzymes that share 4^th^ level E.C. with human essential enzymes. We used available CRISPR gene essentiality data in various human cell lines (Wang et al., 2015) to identify essential genes and enzymes in human cells (Table S5). We further identified essential metabolites in human cells (Table S5) using the latest human genome-scale metabolic model (Robinson et al., 2020) and the metabolic information associated to the essential human genes. Subtracting essential parasite and human enzymes resulted in the analysis of 32 essential *Plasmodium* enzymes catalyzing 68 metabolites and 157 unique metabolite-enzyme pairs in the parasite (Table S6).

### Identification of drugs to target malaria and minimize side effects on human cells

Those molecules that themselves and their downstream products cannot act as inhibitors of essential metabolic enzymes in the human host cell while they can target essential *Plasmodium* enzymes are attractive antimalarial candidates.

We first used NICEdrug.ch to look for small molecules that share reactive site with the 32 essential *Plasmodium* enzymes and they have good similarity score in reactive site neighborhood to native substrates of essential enzymes of parasite, i.e. NICEdrug score above 0.5 (Table S6). We also identified prodrugs that might lead to downstream products with similar reactive site and neighborhood (NICEdrug above 0.5) to any of the essential *Plasmodium* metabolites (Table S6). We suggest those drugs and downstream products act as antimetabolites and competitively inhibit the essential enzymes in the parasite. Overall, we identified 516 drugs that directly compete with essential metabolites and 1,164 prodrugs that need to be biochemically modified between one to three times in human cell to render inhibition of essential enzymes.

We next combined information of essential *Plasmodium* and human metabolites to screen further the drug search using NICEdrug.ch. Out of the hypothetical 516 antimalarial candidates, we identified 64 drugs that share reactive site with parasite metabolites (NICEdrug score above 0.5) and not with human metabolites (NICEdrug score below 0.5), making them good candidates for drug design (Table S6).

### Prediction of inhibitors among food based molecules

We used the reactive site-centric fingerprint available in NICEdrug.ch to identify molecules in food that share reactive site with native substrates of human enzymes and hence might inhibit those enzymes. We retrieved the total set of 80,000 compounds from FooDB (Scalbert et al., 2011), and treated them as input molecules into the NICEdrug pipeline (Figure 1) to identify reactive sites and evaluate their biochemistry, as done for all molecules in NICEdrug.ch.

### Identification of small molecules to target COVID-19

A recent study reported 332 host factors of SARS-CoV-2 (Gordon et al., 2020). Out of the 332 proteins, 97 have catalytic function and EC number assigned, and are potential targets of small molecules. We evaluated the druggability of these 97 enzymes using NICEdrug.ch.

To generate a druggability report, NICEdrug.ch first gathers the metabolic reactions associated with the protein EC numbers. NICEdrug.ch uses 11 databases (including HMR, MetaCyc, KEGG, MetaNetX, Reactome, Rhea, Model SEED, BKMS, BiGG models and Brenda) as source of metabolic reactions. All these databases involve a total of 60k unique metabolic reactions.

Out of the 97 host factor enzymes, we identified 22 enzymes that are linked to fully-defined metabolic reactions. Fully-defined metabolic reactions fulfill three criteria. (1) There is a secondary structure available for all the reaction participants, which means there are available mol files. (2) There is a fully defined molecular structure for all the reaction participants, which means molecules with unspecified R chains are discarded. (3) There is a BNICE.ch enzymatic reaction rule assigned to the reaction (Table S7).

NICEdrug.ch identified 22 host factor enzymes with 24 unique linked EC numbers and 145 unique fully defined reactions. NICEdrug.ch extracts the metabolites participating in these reactions and identifies their reactive site for a reactive-site centric similarity evaluation against a list of molecules. To this end, NICEdrug.ch reports the list of molecules ranked based on the NICEdrug score. The molecule with the highest NICEdrug score shares the highest reactive site-centric similarity with the native substrate of the target enzyme (Table S7).

We found 1,301 molecules that show NICEdrug score above 0.5 with respect to substrates of the 22 SARS-CoV-2 hijacked enzymes (Table S7). Out of 1,301 molecules, 465 are drugs cataloged in DrugBank, KEGG drugs or ChEMBL databases, 712 are active molecules one step away of 1,419 prodrugs, and 402 are food molecules (Table S7).

To better understand the classes of drugs or food molecules, we classified drugs based on their KEGG drug groups (Dgroups) and food molecules based on their food source. Out of 465 drugs identified, 43 drugs are assigned to 55 different Dgroups and 402 food molecules belong to 74 different food sources (Table S7).

## Supplemental table titles and legends

**Table S1. Information of human metabolism considered in this study, related to Figures 2, 3, 4, 5, and 6.**

**(A)** List of cofactors, **(B)** list of metabolites, and **(C)** list of E.C. numbers considered in BNICE.ch for the generation of reactions in the analysis of drug metabolism in a human cell (Materials and Methods).

**Table S2. Metabolic neighborhood of 5-FU, related to Figures 2, 3, and 4.**

**(1)** List of compounds in the 5-FU metabolic neighborhood including up to four reactions or steps away. **(2)** Description of reactions in the 5-FU metabolic neighborhood including up to four reactions or steps away.

**Table S3. NICEdrug score between all molecules with reactive site of statins in NICEdrug.ch, related to Figure 5.**

Matrix of NICEdrug score between each pair of the whole set of 254 molecules in NICEdrug.ch with reactive site of statins.

**Table S4. Description of nine drugs candidates for repurposing to replace statins based on NICEdrug.ch, related to Figure 5.**

These drugs can act as competitive inhibitors of HMG-CoA reductase like statins.

**Table S5. Essential genes or enzymes and linked metabolites in liver-stage *Plasmodium* and a human cell, related to Figure 6.**

**(A)** List of essential genes and associated reactions in liver-stage *Plasmodium*, as obtained from the study (Stanway et al., 2019) **(B)** List of essential genes and associated reactions in a human cell, as obtained from the study (Wang et al., 2015) **(C)** List of metabolites linked to essential genes in liver-stage *Plasmodium*. **(D)** List of metabolites linked to essential genes in a human cell.

**Table S6. Description of drugs, prodrugs, metabolites and enzymes analyzed in the study of malaria, related to Figure 6**

**(A)** NICEdrug druggability analysis of essential genes or enzymes in liver-stage *Plasmodium*: all drugs sharing reactive-site centric similarity with the *Plasmodium* metabolites and comparison with human metabolites. **(B)** NICEdrug druggability analysis of essential genes or enzymes in liver-stage *Plasmodium*: all prodrugs (up to three steps away of 346 drugs) sharing reactive-site centric similarity with the *Plasmodium* metabolites and comparison with human metabolites. **(C)** Description of drugs and prodrugs identified in the malaria analysis with NICEdrug.ch and validated in the study by (Antonova-Koch et al., 2018) along with their similar *Plasmodium* metabolite and human metabolite.

**Table S7. Hijacked human enzymes by SARS-CoV-2, and drugs and food-based compounds that can inhibit them based on the NICEdrug score, related to Figure 7.**

**(A)** Hijacked human proteins by SARS-CoV-2 as identified by (Gordon et al., 2020) with an annotated enzymatic function (E.C. number), also called here “SARS-CoV-2 hijacked enzymes”. **(B)** NICEdrug druggability report for SARS-CoV-2 hijacked enzymes including all NICEdrug small molecules. **(C)** Best candidate drugs against COVID-19: NICEdrug druggability report for SARS-CoV-2 hijacked enzymes including drugs with NICEdrug score above 0.5 compared to the native human substrate. **(D)** Summary of NICEdrug best candidate drugs against COVID-19 and their classification according to the drug category in the KEGG database. **(E)** NICEdrug druggability report of SARS-CoV-2 hijacked enzymes including prodrugs (up to three steps away of any NICEdrug small molecule) with NICEdrug score above 0.5 compared to the native human substrate. **(F)** Best candidate food-based molecules against COVID-19: NICEdrug druggability report of SARS-CoV-2 hijacked enzymes including food-based molecules with NICEdrug score above 0.5 compared to the native human substrate. **(G)** Summary of the NICEdrug best candidate food-based molecules against COVID-19 and their classification according to the fooDB source.

**Table S8. NICEdrug analysis of inhibitory mechanisms of currently used anti SARS-CoV-2 drugs, related to Figure 7.**

**(A)** All drug molecules and **(B)** prodrugs in NICEdrug.ch sharing reactive site with the native substrates of the human enzyme HDAC2 and their NICEdrug score with this substrate. **(C)** All molecules cataloged in fooDB sharing reactive site with the native substrates of the human enzyme HDAC2 and their NICEdrug score with this substrate. **(D)** All drug molecules and **(E)** prodrug molecules in NICEdrug.ch sharing reactive site with the native substrates of the human enzyme ACE2 and their NICEdrug score with this substrate. **(F)** All molecules cataloged in fooDB sharing reactive site with the native substrates of the human enzyme ACE2 and their NICEdrug score with this substrate. **(G)** All molecules in NICEdrug.ch or cataloged in fooDB sharing reactive site with the native substrates of the human enzyme DNA-directed RNA polymerase and their NICEdrug score with this substrate.

